# m^6^A demethylase ALKBH5 promotes tumor cell proliferation by destabilizing IGF2BPs target genes and worsens the prognosis of patients with non-small cell lung cancer

**DOI:** 10.1101/2021.07.06.451216

**Authors:** Kazuo Tsuchiya, Katsuhiro Yoshimura, Yuji Iwashita, Yusuke Inoue, Tsutomu Ohta, Hirofumi Watanabe, Hidetaka Yamada, Akikazu Kawase, Masayuki Tanahashi, Hiroshi Ogawa, Kazuhito Funai, Kazuya Shinmura, Takafumi Suda, Haruhiko Sugimura

**Affiliations:** Department of Tumor Pathology, Hamamatsu University School of Medicine, Hamamatsu, Japan; Second Division, Department of Internal Medicine, Hamamatsu University School of Medicine, Hamamatsu, Japan; Department of Physical Therapy, Faculty of Health and Medical Sciences, Tokoha University, Hamamatsu, Japan; First Department of Surgery, Hamamatsu University School of Medicine, Hamamatsu, Japan; Division of Thoracic Surgery, Respiratory Disease Center, Seirei Mikatahara General Hospital, Hamamatsu, Japan; Department of Pathology, Seirei Mikatahara General Hospital, Hamamatsu, Japan

**Keywords:** Non-small cell lung cancer (NSCLC), m^6^A, ALKBH5, IGF2BPs, CDKN1A (p21), TIMP3

## Abstract

The modification of *N*^6^-methyladenosine (m^6^A) in RNA and its eraser ALKBH5, an m^6^A demethylase, play important roles across various steps of human carcinogenesis. However, the involvement of ALKBH5 in non-small cell lung cancer (NSCLC) development remains to be completely elucidated. The current study revealed that the expression of ALKBH5 were increased in NSCLC and increased expression of ALKBH5 worsened the prognosis of patients with NSCLC. *In vitro* study revealed that ALKBH5 knockdown suppressed cell proliferation ability of PC9 and A549 cells as well as promoted G1 arrest and increased the number of apoptotic cells. Furthermore, ALKBH5 overexpression increased the cell proliferation ability of the immortalized cell lines. Microarray analysis and western blotting revealed that the expression of CDKN1A or TIMP3 were increased by ALKBH5 knockdown. These alterations were offset by a double knockdown of both ALKBH5 and one of the IGF2BPs. The decline of mRNAs was, at least partly, owing to the destabilization of these mRNAs by one of the IGF2BPs. In conclusions, the ALKBH5–IGF2BPs axis promotes cell proliferation and tumorigenicity, which in turn causes the unfavorable prognosis of NSCLC.

## Background

Lung cancer, the incidence of which has continued to increase annually, remains the most frequently diagnosed cancer and leading cause of cancer-related death worldwide [1]. Considering the rapid improvement in the treatment of lung cancer, particularly non-small cell lung cancer (NSCLC), physicians now have several options of personalized treatments targeting driver genes, such as EGFR mutations, ALK rearrangements, ROS1 rearrangements, and BRAF mutations or combination therapies comprising immunotherapy and anticancer drugs [2–6]. However, despite the current advancements in precision medicine, NSCLC still exhibits poor long-term prognosis and high mortality rates owing to the rapid growth, metastasis, and infiltration of cancer. Therefore, identifying effective therapeutic targets that inhibit such malignant behaviors of NSCLC is urgently needed.

*N*^6^-methyladenosine (m^6^A), the most prevalent internal messenger RNA (mRNA) modification, controls various mRNA functions. The m^6^A sites, which are widely distributed around the stop codons and 3′ untranslated regions (UTRs) of mRNAs, presumably exist in precursor mRNAs [7]. Recent m^6^A transcriptome analysis revealed that m^6^A is predominantly present in the RRACU (R = A/G) consensus motif of mammals [8]. m^6^A is reversibly catalyzed by a methyltransferase complex (writer) and demethylase (eraser). Accordingly, the methyltransferase complex comprises methyltransferase-like 3 and 14 (METTL3 and METTL14) with their cofactors Wilms tumor 1-associated protein (WTAP), VIRMA (KIAA1429), and RNA-binding motif protein 15 (RBM15) [9–12]. Further, fat mass and obesity-related protein (FTO) and AlkB homolog 5 (ALKBH5) have been identified as two eukaryotic demethylases that oxidatively demethylate m^6^A with α-ketoglutarate as a substrate and Fe (II) as a coenzyme [13, 14]. FTO is also involved in the demethylation of *N*^6^,2 -*O*-dimethyladenosine (m^6^A_m_), whereas other than m6A, there is no other RNA that is demethylated by ALKBH5 [15, 16]. Moreover, m^6^A-binding protein (reader protein), which recognizes m^6^A, is involved in numerous biological processes in an m^6^A-dependent manner. For instance, YT521-B homology (YTH) domain containing 1 (YTHDC1) and YTHDC2 promote alternative splicing and mRNA export from the nucleus to the cytoplasm [17, 18]; heterogeneous nuclear ribonucleoprotein G (HNRNPG) alters RNA structures via RNA–protein interaction [19, 20]; YTH domain family1 (YTHDF1), YTHDF3, METTL3, and eukaryotic initiation factor3 (eIF3) regulate translation efficiency [21–24]; and insulin-like growth factor 2 mRNA-binding proteins (IGF2BPs), YTHDF2, YTHDF3, and YTHDC2 alter mRNA stability [24–27].

Over the past few years, several researchers investigating the role of m^6^A eraser in malignant tumors have revealed that m^6^A eraser proteins play a critical role in oncogenesis. A number of previous studies have demonstrated that ALKBH5 exerts a cancer-promoting effect in glioblastoma, osteosarcoma, colon cancer, ovarian cancer, esophageal squamous cell carcinoma, endometrial cancer, and renal cell carcinoma [28–34]. In contrast, ALKBH5 has been reported to play a tumor-suppressing effect in hepatocellular carcinoma and pancreatic cancer [35, 36]. Several studies on lung cancer have shown that FTO plays a cancer-promoting role through m^6^A modification in lung squamous cell carcinoma and adenocarcinoma [37–39], whereas ALKBH5 inhibits NSCLC tumorigenesis by reducing YTHDFs-mediated YAP expression [40]. Conversely, ALKBH5 had also been found to promote NSCLC progression by regulating TIMP3 stability [41]. Thus, the precise role of ALKBH5 in NSCLC tumorigenesis across various conditions deserves further investigation.

Cell type and cell environment (e.g., during hypoxic conditions) as well as the m^6^A target gene and its recognition protein (reader) have been found to affect RNA metabolism caused by ALKBH5 perturbation [42, 43]. Therefore, m^6^A-mediated gene expression regulated by ALKBH5 could result in various consequences in cancer cells depending on the surrounding environment and other factors. Several studies regarding m^6^A have focused on specific genes in the specific contexts, with their results showing that m^6^A is involved in the mechanisms through which these specific genes are regulated. However, in actual human cancers, ALKBH5 catalyzes specific m^6^A of numerous genes, which simultaneously alters several RNA and protein expressions through RNA recognition by reader proteins, consequently causing numerous interactions between them *in vivo*. As such, systematically clarifying the association between m^6^A modification and cancer development across each clinical and pathological setting is important. Furthermore, elucidating the significance of m^6^A modification by ALKBH5 may facilitate the clinical usage of such molecules as therapeutic targets. Therefore, the current study aimed to examine the role of m^6^A demethylase in NSCLC focusing on ALKBH5 and determine its association with downstream targets, including “readers” and “target genes.”

## Methods

### Immunohistochemistry

Resected NSCLC samples from Hamamatsu University School of Medicine and Seirei Mikatahara General Hospital were collected and named as the HUSM cohort. Tissue microarray (TMA) sections were analyzed using immunohistochemistry (IHC) as previously described [44]. Cores of insufficient quality or quantity were excluded from analysis. Antibodies for ALKBH5 (HPA007196, Atlas Antibodies, Stockholm, Sweden) and FTO (Ab124892, Abcam, Cambridge, UK) were diluted at 1:400, whereas those specific for EGFR E746-A750 deletion (#2085, D6B6, Cell Signaling Technology [CST], Danvers, MA, USA) and EGFR L858R mutant (#3197, 43B2, CST) were diluted at 1:100, followed by incubation at room temperature for 0.5 h. Protein expression levels were then assessed using the H-score, which was calculated by multiplying the percentage of stained tumor area (0%–100%) by the staining intensity (scored on a scale of 0–3) to yield a value ranging from 0 to 300.

### Analysis of publicly available datasets

We used the lung cancer database in the Kaplan–Meier plotter (http://kmplot.com/analysis/index.php?p=service&cancer=lung) to analyze the association between prognosis and ALKBH5 and FTO mRNA expression in NSCLC cohorts. Data were downloaded on December 10, 2020. Kaplan–Meier curves for overall survival (OS) were generated and stratified according to the median expression of each mRNA. To assess the mRNA expression of ALKBH5 and FTO, data from the Cancer Genome Atlas (TCGA) (NSCLC, Provisional) were downloaded from cBioPortal (http://www.cbioportal.org/) on November 11, 2019. Expression data were obtained in the form of RNA-seq by Expectation Maximization (RSEM).

### Immunofluorescence analysis

Cells grown on coverslips were fixed with 4% paraformaldehyde and permeabilized with 0.1% Triton X-100. After blocking with 5% bovine serum albumin in PBS (−) at room temperature for 1 h, the cells were probed with primary antibodies against ALKBH5 (HPA007196, Atlas Antibodies) and then incubated with a Goat anti-Rabbit IgG (H + L) Cross-Adsorbed Secondary Antibody, Alexa Fluor 546 (#A-11010, Thermo Fisher Scientific, Waltham, MA, USA). Nuclei were stained with ProLong® Gold Antifade Reagent with DAPI (#8961, CST), after which the cells were imaged via fluorescence microscopy using z-stack image reconstructions (BZ-9000; Keyence, Osaka, Japan).

### Cell lines and transient knockdown with siRNA

The human lung cancer cell lines H1299, H460, H2087, A549, ABC1, and H358 and the human immortalized cell lines BEAS2B and HEK293 were obtained from Health Science Research Resources Bank (Osaka, Japan) or the American Type Culture Collection (Manassas, VA, USA). PC3 and PC9 lung cancer cells were purchased from the Japanese Collection of Research Bioresources Cell Bank (Osaka, Japan) and RIKEN BioResource Center (Tsukuba, Japan), respectively, whereas ACC-LC176 cells were a kind gift from Dr. Takashi Takahashi (Nagoya University). RERF-LC–MS, HLC-1, and LC-2/ad were kind gifts from Dr. Toshiro Niki (Tokyo University). Lung cancer cell lines were cultured in RPMI1640 medium (R8758, Thermo Fisher Scientific), whereas HEK293 cells were cultured in DMEM (D5796 MERCK, Darmstadt, Germany) containing 10% (vol./vol.) fetal bovine serum (FBS), 100 IU/mL penicillin G, and 100 µg/mL streptomycin. LHC9 (12680013, Thermo Fisher Scientific) was also used as a medium for BEAS2B cells. Cells were maintained in a 5% CO_2_ and 95% air incubator at 37□. Silencer Select Pre-designed siRNA for ALKBH5 (siALKBH5: s29743, s29744, s29745, Invitrogen), FTO (siFTO: s28147, s28148, s28149, Invitrogen), IGF2BP1(siIGF2BP1: s20916, s20917, Invitrogen, Carlsbad, CA, USA), IGF2BP2 (siIGF2BP2: s20922, s20923, Invitrogen), IGF2BP3(siIGF2BP3: s20919, s20920, Invitrogen), and Silencer Select Negative control (siNC: 4390843, Invitrogen) were purchased for transient knockdown. More than two different sequences were used for one target gene to minimize off-target effects. Cells were cultured for 24 h before transfection, after which they were transfected with 15 nM of final siRNA concentrations using Opti-MEM (31985070, Gibco, Dublin, Ireland) and Lipofectamine® 2000 (11668019, ThermoFisher). The cells were then used for further assays at 48–96 h after transfection. When no siRNA sample number was available, siRNA no. 1 (#1) and siRNA no. 3 (#3) were pooled for ALKBH5 unless otherwise specified. siIGF2BP1, siIGF2BP2, and siIGF2BP3 were pooled for all transfections.

### Generation of Retro-X Tet-On inducible cell lines overexpressing ALKBH5

The retroviral plasmid pRetroX-TetOne puro (634307, Clontech, Mountain View, CA, USA) was amplified using NEB Stable competent *Escherichia coli* (high efficiency) (C3040H, NEW ENGLAND BioLabs, Ipswich, MA, USA). The full-length ALKBH5 sequence (NM_017758), which was confirmed using Sanger sequencing, was subcloned into pRetroX-TetOne puro vector using EcoRI and BglII restriction sites (pRetroX-TetOne puro-ALKBH5). Retroviral supernatants were produced using the GP2-293 packaging cell line (Clontech), in which pRetroX-TetOne puro empty vector or pRetroX-TetOne puro-ALKBH5 were each cotransfected with the envelope vector VSV-G using Xfect transfection reagent (Clontech). BEAS2B, HEK293, PC9, and A549 cells were transfected for 24 h using 4 µg/mL polybrene (H9268, Sigma-Aldrich, St. Louis, MO, USA). Puromycin selection (0.5–1.5 µg/mL) began 48 h after transfection and lasted for 3 days until all non-transfected cells had died. In subsequent experiments, Retro-X cells were induced with 0.1–100 ng/mL of doxycycline (DOX) diluted in culture media upon cell seeding for 24–96 h. Cells transfected with pRetroX-TetOne puro-ALKBH5 with 100 ng/mL DOX were designated as ALKBH5-overexpressed (OE) cells, whereas those infected without DOX were designated as negative control (NC) unless otherwise noted.

### RNA isolation and quantitative-polymerase chain reaction (qPCR)

Total RNA was extracted using the RNeasy Plus Mini Kit (#74136, QIAGEN, Hilden, Germany) according to the manufacturer’s instructions, with the total RNA concentration calculated using Nanodrop (NanoDrop1000, Thermo Fisher Scientific). cDNA was synthesized from 1 µg of total RNA using the ReverTra Ace qPCR RT Master Mix (FSQ-201, TOYOBO) according to the manufacturer’s instructions. qPCR reactions were performed on a Step One Plus Real-Time PCR System (Applied Biosystems, Thermo Fisher Scientific) using the THUNDERBIRD qPCR Mix (QPS-201, TOYOBO, Osaka, Japan). The relative RNA expression levels were calculated using the ΔΔCt method, with the levels normalized to glyceraldehyde 3-phosphate dehydrogenase (GAPDH) mRNA. All amplicons were confirmed as a single product using agarose gel visualization and/or melting curve analysis. The applied primer sequences are listed in Table S1.

### Protein isolation and western blotting

Total protein lysates were extracted from whole cells using 1× sodium dodecyl sulfate (SDS) sample buffer. The Pierce BCA Protein Assay Kit (Cat#23225, Thermo Fisher Scientific) was used to determine the protein concentration. All proteins were separated using SDS-polyacrylamide gel electrophoresis and transferred to PVDF Blotting Membrane (P 0.45, A29532146, GE healthcare Life science, Chicago, IL, USA) using the Trans-Blot Turbo Cassette (Bio-Rad, Hercules, CA, USA). Blocking One (03953, Nacalai, Kyoto, Japan) or 5% skimmed milk were used for blocking. Primary antibodies for ALKBH5 (1:1000 dilution, HPA007196; Atlas Antibodies), FTO (1:1000 dilution, Ab124892; Abcam), IGF2BP1 (1:1000 dilution, 22803-1-AP; Proteintech), IGF2BP2 (1:1000 dilution, 11601-1-AP; Proteintech), IGF2BP3 (1:2000 dilution, 14642-1-AP; Proteintech), TIMP3 (1:3000 dilution, Ab39184; Abcam), p21 (1:1000 dilution, A1483; ABclonal, Woburn, MA, USA), E2F1 (1:500 dilution, A2067; ABclonal), CCNG2 (0.2μg/mL, Ab251826; Abcam), p53 (1:200 dilution, Sc-126; SANTA CRUZ BIO TECHNOLOGY, Dallas, TX, USA), and GAPDH (1:1000 dilution, Ab8245; Abcam) were incubated for overnight at 4□. Secondary antibodies for rabbit (1:20000 dilution, NA9340; GE healthcare Life science) or mouse (1:20000 dilution, NA9310; GE Healthcare Life Science) were incubated at room temperature with 1%–5% skimmed milk for 1 h. Enhanced chemiluminescence (Pierce ECL Plus Substrate or West Atto Ultimate Sensitivity Substrate, Thermo Fisher Scientific) was used to visualize the protein bands using ChemiDocTouch (Bio-Rad).

### Cell viability assay

Cells were seeded into 96-well plates with 3000 cells per well after 48 h of knockdown or overexpression. Cell proliferation was monitored using Cell Counting Kit-8 (CCK-8; Dojindo, Kumamoto, Japan) according to the manufacturer’s protocol. Thereafter, the cells were incubated with 10% CCK-8 for 1 h, followed by absorbance assessment at 450 nm in each well via spectrophotometry (Synergy HT, BioTek, Winooski, VT, USA) every 24 h.

### Transwell migration assay

Cell migration was evaluated using a 24-well plate with cell culture inserts (353097, Falcon, Mexico City, Mexico) containing a filter with 8 μm-diameter pores. Briefly, after serum starvation for 24 h with 0.1% FBS-containing RPMI1640 medium, 1 × 10^5^ cells resuspended in 500 μL of RPMI1640 medium (Gibco) were seeded into the upper chamber, after which RPMI1640 medium containing 10% FBS was placed in the lower compartment of the chamber. After incubation for 16 h, the upper surface of the membrane was wiped with a cotton-tipped applicator to remove non-migrating cells, whereas the migrating cells on the lower surface were fixed with cold methanol and stained with 0.5% crystal violet. Migrating cells were automatically counted in three random microscopic fields using the Hybrid Cell Count software (BZ-□ Analyzer, Keyence, Osaka, Japan).

### Wound-healing assays

To assess cell migration, 2 × 10^5^ cells were seeded into 6-well plates. Thereafter, cells were incubated in 5% CO_2_ at 37□ for 48□h and an additional 24 h with 0.1% FBS-containing RPMI1640 medium. A wound was scratched into the cells using a 200-μL plastic tip and washed with PBS (−). The cells were then incubated in RPMI1640 containing 10% FBS. The relative distance of the scratches was observed under an optical microscope (IX53, Olympus, Tokyo, Japan) at 3–6 time points after wounding and assessed using the Image J software.

### Cell cycle assay and apoptosis assay

Cell Cycle Assay Solution Blue (C549, Dojindo) was used to measure the cell cycle according to the manufacturer’s instructions. Briefly, treated cells were synchronized at the G1 phase through serum starvation with 0.1% FBS-containing medium for 48 h. At 24 h after the release of serum starvation, the treated cells were collected, washed with PBS (−), and incubated with 5 μL cell cycle assay solution for 15 min at 37□. Thereafter, DNA content was determined based on staining intensity using a Gallios flow cytometer (Beckman Coulter, Miami, FL, USA). The Annexin V-FITC Apoptosis Detection Kit (15342-54 Nacalai) was used to detect apoptosis by measuring annexin V and propidium iodide (PI)-positive cells following the manufacturer’s instructions. Briefly, cells were incubated for 96 h after siRNA transfection. To induce apoptosis, the cells were exposed to either 7.5 μM of gefitinib (078-06561, FUJIFILM) or 10 μM of cisplatin (P4394, Sigma-Aldrich) alone for 48 h after siRNA transfection. The treated cells were collected, washed with PBS (−), and incubated with 5 μL of annexin V-FITC solution and 5 μL of PI solution for 15 min. Thereafter, apoptotic cells were determined using a Gallios flow cytometer. Results were analyzed using the FlowJo software (Becton, Dickinson, Franklin Lakes, NJ, USA), after which the extent of apoptosis and cell cycle distribution were determined.

### RNA stability assay

Cancer cells were incubated for 48 h after siRNA transfection. Cells were treated with actinomycin D at a final concentration of 5 μg/mL. Total RNA was extracted at 0, 2, 4, and 6 h after adding actinomycin D. The remaining CDKN1A and TIMP3 mRNA was measured through quantitative real-time PCR and normalized to RPL32 mRNA, which has a half-life of 25 h.

### Quantitative analysis of global m^6^A levels using liquid chromatography–mass spectrometry/mass spectrometry (LC–MS/MS)

PolyA-enriched RNA was extracted using PolyATract mRNA isolation systems (#Z5310 Promega, Madison, WI, USA) according to the manufacturer’s instructions. PolyA-enriched RNA concentration was calculated using Qubit 2.0. The polyA-enriched RNA was enzymatically hydrolyzed using 8-OHdG Assay Preparation Reagent Set (292-67801, FUJIFILM Wako Pure Chemical Corporation, Tokyo, Japan). Technically, 100 ng of polyA-enriched RNA was digested using 5.7 μL of acetic acid buffer and 3 μL of Nuclease P1 included in the 45-μL sample containing nuclease-free water at 37□ for 30 min, followed by incubation with 6 μL of Tris Buffer and 0.3 μL of alkaline phosphatase at 37□ for 30 min. After digestion, the sample was centrifuged at 14,000 *g* and 4□ for 20 min using a Nanosep 3K Omega centrifugal device (Pall Corporation, Port Washington, NY, USA) according to a previously published method [45].

As an internal standard, *N*^6^-methyladenosine-d3 (m^6^A-d3; M275897, Toronto Research Chemicals, Toronto, Canada), which is a stable isotope of *N*^6^-methyladenosine labeled with three deuterium atoms on the *N*^6^-methyl group, was added to the nucleosides obtained via digestion of polyA-enriched RNA. These nucleosides were separated using an Acquity UPLC HSS T3 column (2.1 mm × 100 mm; Waters, Milford, CT, USA) with 0.1% (vol./vol.) formic acid in water as mobile phase A and methanol as mobile phase B at a flow rate of 200 μL/min in a linear gradient elution of 5%–60% B from 0 to 7 min. Standard compounds of adenosine (A; A9251, Sigma-Aldrich), *N*^6^-methyladenosine (m^6^A; A170736, Sigma-Aldrich), and m^6^A-d3 were used to confirm the nucleoside-to-base ion mass transitions of 268.1–136.4 (A), 282.2–150.2 (m^6^A), and 285.2–153.2 (m^6^A-d3). Peak areas of A, m^6^A, and m^6^A-d3 in the nucleosides digested from polyA-enriched RNA were calculated using the column retention time of the standard compounds using Analyst 1.6.1 software (AB SCIEX, Foster City, CA, USA). The m^6^A level was quantified as the ratio of m^6^A to A or m^6^A-d3 based on the calibrated concentrations.

### Microarray analysis of differentially expressed genes

Total RNA was extracted from ALKBH5-knockdown or control PC9 cells 96 h after transfection. RNA samples were used for global gene expression profiling on human Clariom S Assay microarrays (Thermo Fisher Scientific, Wilmington, DE, USA), which include 24351 genes. All microarray analyses were entrusted to Filgen Inc. (Aichi, Japan). A total RNA quality control check was performed using a NanoDrop ND-1000 (Thermo Scientific) and an Agilent 2100 Bioanalyzer. Using the Gene Chip TM WT PLUS Reagent Kit, fragmented and labeled cDNA samples were prepared from 250 ng of total RNA according to the manufacturer’s instructions (Gene Chip TM WT PLUS Reagent Kit User Manual). Thereafter, Thereafter, 100 L of hybridization solution was prepared using 73 μL of Hybridization Master Mix and 2.3 μg of fragmented and labeled cDNA. The array was incubated using the Gene Chip TM Hybridization Oven 645 at 45□ for 16 h (60 rpm). The array was cleaned using the Gene Chip TM Fluidics Station 450 and scanned using the Gene Chip TM Scanner 3000 7G according to the manufacturer’s instructions [Gene Chip TM Command Console (AGCC) 4.0 User Manual]. The Microarray Data Analysis Tool version 3.2 (Filgen, Aichi, Japan) was used for data normalization and subsequent processing. Differentially expressed mRNAs were identified using a set cutoff (fold change > 1.5 or < 0.67; *P* < 0.01). Gene set enrichment analysis (GSEA) was performed to examine the gene set regulated by ALKBH5 knockdown (http://software.broadinstitute.org/gsea/omdex.jsp). For analysis, the false discovery rate (FDR) based on gene set permutation was used. Microarray data has been deposited in the Gene Expression Omnibus (GEO) at the National Center for Biotechnology Information (NCBI) (accession number GSE165453).

### Epitranscriptomic microarray analysis

Unfragmented total RNA was extracted from ALKBH5-knockdown or control PC9 cells at 96 h after transfection and quantified using the NanoDrop ND-1000. RNA samples were used for global m^6^A expression profiling on an Arraystar Human mRNA&lncRNA Epitranscriptomic Microarray (8 × 60 K; Arraystar), which includes 44,122 protein-coding mRNAs and 12,496 long non-coding RNAs. Microarray analyses were entrusted to Arraystar Inc. (Rockville, MD, USA). Sample preparation and microarray hybridization were performed based on Arraystar’s standard protocols. Briefly, total RNAs were immunoprecipitated with an anti-m^6^A antibody (Synaptic Systems, 202003). The “immunoprecipitated (IP)” and “supernatant (Sup)” RNAs were labeled with Cy5 and Cy3, respectively, as cRNAs in separate reactions using the Arraystar Super RNA Labeling Kit. The cRNAs were combined and hybridized onto Arraystar Human mRNA&lncRNA Epitranscriptomic Microarray (8 × 60 K, Arraystar). After washing the slides, the arrays were scanned in two-color channels using an Agilent Scanner G2505C. Agilent Feature Extraction software (version 11.0.1.1) was used to analyze acquired array images. Raw intensities of IP (Cy5-labeled) and Sup (Cy3-labeled) were normalized with an average of log2-scaled Spike-in RNA intensities. The “m^6^A methylation level” was calculated to determine the percentage of modification based on the IP (Cy5-labeled) and Sup (Cy3-labeled) normalized intensities. “m^6^A quantity” was calculated to determine the amount of m^6^A methylation based on the IP (Cy5-labeled) normalized intensities. Differentially m^6^A-methylated RNAs between both comparison groups were identified by filtering with a fold change of >1.5 or <0.67 (*P* < 0.01) through the unpaired *t*-test. Microarray data had been deposited in the GEO at the NCBI (accession number GSE165454).

### qPCR for methylated RNA immunoprecipitation (MeRIP) with m^6^A antibody

ALKBH5-knockdown or control lung cancer cells were used for methylated RNA immunoprecipitation assay. The Magna MeRIP m^6^A kit (catalog no.17-10499, Millipore, Burlington, MA, USA) was used according to the manufacturer’s protocol. Briefly, the polyA-enriched RNA was fragmented into 100–200 nucleotides incubated with RNA fragmentation buffer for 55 s (CS220011, Millipore). The size of polyA-enriched RNA fragments was optimized using the Agilent 4200 TapeStation (Agilent technologies, Santa Clara, CA, USA). We used 0.5 μg of fragmented polyA-enriched RNA as input control and 5 μg of fragmented polyA-enriched RNA for m^6^A mRNA immunoprecipitation, followed by incubation with m^6^A antibody (MABE1006, Millipore)- or mouse IgG-conjugated Protein A/G Magnetic Beads in 500 μL 1× IP buffer supplemented with RNase inhibitors at 4□ overnight. Methylated RNAs were immunoprecipitated with beads, eluted via competition with free m^6^A, and purified using the RNeasy kit (Qiagen). Moreover, modification of m^6^A toward particular genes was determined using qPCR analysis with specific primers [primers for the positive control region (stop codon, EEF1A1+) or NC region (exon 5, EEF1A1−) of human EEF1A1 was included in the Magna MeRIP m6A kit]. To design primers for MeRIP qPCR, m^6^A sites of specific genes were predicted using the sequence-based RNA adenosine methylation site predictor algorithm (http://www.cuilab.cn/sramp) [46]. We focused on the potential m^6^A sites in the 3′ UTRs near the stop codon and designed primers to ensure that the target sequence were present in these sites with a limited length of 120□nt. Self-designed primers for MeRIP qPCR are listed in Table S1.

### Statistical analysis

Discrete variables were expressed as numbers (percentages), whereas continuous variables were expressed as means ± standard deviations (SDs) unless otherwise specified. The Mann–Whitney *U* test was used to compare continuous individual samples, whereas Student’s *t*-test was applied to compare continuous experimental data. Fisher’s exact test for independence was used to compare categorical data between groups. The Wilcoxon matched-pairs signed-rank test was used to compare two corresponding groups. Spearman’s correlation coefficient was used for correlation analysis. Kaplan–Meier curves with log-rank tests were used to analyze survival. Accordingly, OS was defined as the duration from baseline to the date of death, whereas recurrence-free survival (RFS) was defined as the duration from baseline to the recurrence date. Univariate and multivariate Cox proportional hazards models were applied to generate hazard ratios (HRs) for death while adjusting for other potential confounding factors. Cell proliferation and RNA stability assays were analyzed using two-way analysis of variance. Statistical analyses were performed using GraphPad Prism Version 8 (GraphPad Software, San Diego, CA, USA) and EZR software (Saitama Medical Center, Jichi Medical University, Saitama, Japan), with *P* values of <0.05 indicating statistical significance.

## Results

### High ALKBH5 expression was associated with a worse prognosis in patients with NSCLC

To investigate the impact of ALKBH5 and FTO in NSCLC, we examined the mRNA expression levels of ALKBH5 and FTO in non-cancerous lung tissues and NSCLC tissues using TCGA data. Accordingly, our results showed no significant difference in ALKBH5 mRNA expression between non-cancerous and cancerous tissues. By contrast, our findings showed that NSCLC had a significantly lower FTO mRNA expression than non-cancerous tissues (Figure 1a). We subsequently investigated the protein expression levels of ALKBH5 and FTO in non-cancerous lung alveolar tissue and corresponding NSCLC tissues using TMA of patient samples. Furthermore, our results showed that cancerous tissues had significantly higher H-scores for ALKBH5 and FTO than non-cancerous tissues (Figure 1b). ALKBH5 and FTO expression were evaluated in immortalized bronchial epithelial cells (BEAS2B) and lung cancer cell lines. Consequently, qPCR analysis demonstrated that ALKBH5 mRNA expression was higher in lung cancer cell lines except LC-2/ad and RERF-LC–MS, whereas FTO mRNA expression was lower in lung cancer cell lines except HLC-1, ABC1, and PC3 (Figure 1c). Western blot analysis demonstrated that ALKBH5 and FTO were endogenously expressed in all lung cancer cell lines except HLC-1 (Figure 1d). IHC analysis showed that ALKBH5 and FTO were mainly localized in the nucleus of the cells (Figure 1e). Furthermore, immunofluorescence analysis showed that ALKBH5 was localized in the nucleus of PC9 cells overexpressing ALKBH5 (Figure 1f). We analyzed the clinical characteristics of 627 NSCLC cases used in IHC of TMA in the context of ALKBH5 or FTO expression in tumors of the HUSM cohort (Table S2). The median age was 68 (range, 23–88) years; 430 (68.6%) patients were male and 184 (29.3%) had never smoked. The tumors were histologically classified as adenocarcinoma (n = 413, 65.9%), squamous cell carcinoma (n = 170, 27.1%), or other histological types (n = 44, 7.0%). A total of 395 (63.0%) patients had stage Ι disease, whereas 127 (20.3%) cases had EGFR mutations. Postoperative adjuvant chemotherapy was prescribed to 258 (41.1%) patients. The median H-score values for ALKBH5 and FTO expression was 110 (0–225, range) and 65 (0–281), respectively, with high ALKBH5 protein expression being correlated with high FTO protein expression (r = 0.41) (Figure S1A). Based on the median value, cases were divided into “high” and “low” expression groups, after which their association with clinical data as well as prognostic significance was examined. Lymph node metastasis, chemotherapy, and EGFR status significantly differed depending on ALKBH5 expression, whereas tumor status, lymph node metastasis, pathological stage, chemotherapy, and EGFR status significantly differed depending on FTO expression. Kaplan–Meier curves showed that patients with high ALKBH5 expression had significantly worse survival than those with low ALKBH5 expression (Figure 1g: log-rank p = 0.0009 for OS; Figure S1B: log-rank p = 0.0008 for RFS). Conversely, Kaplan–Meier curves showed no significant difference in survival between the low and high FTO expression groups (Figure 1g: log-rank p = 0.20 for OS, Figure S1C: log-rank p = 0.07 for RFS). Univariate analysis revealed high ALKBH5 expression as a predictor of unfavorable OS (HR, 1.675; 95% CI, 1.230–2.521). Moreover, multivariate analysis of age, sex, smoking status, histology, pathological stage, and ALKBH5 expression revealed that ALKBH5 expression was an independent prognostic factor associated with unfavorable OS (HR, 1.468; 95% CI, 1.039–2.073) (Table S3). To validate the prognostic value of ALKBH5 and FTO in other cohorts of patients with NSCLC, the lung cancer database in the Kaplan–Meier plotter was used. Accordingly, Kaplan–Meier curves showed that patients with high ALKBH5 expression had a significantly worse survival than those with low ALKBH5 expression (Figure S1D: log-rank p = 0.014 for OS). In contrast, Kaplan–Meier curves showed that patients with high FTO expression had significantly favorable survival compared with those with low FTO expression (log-rank p < 0.0001 for OS) (Figure S1E). These observations suggested that ALKBH5 played a critical role in the poor prognosis of patients with NSCLC.

**Figure 1.**
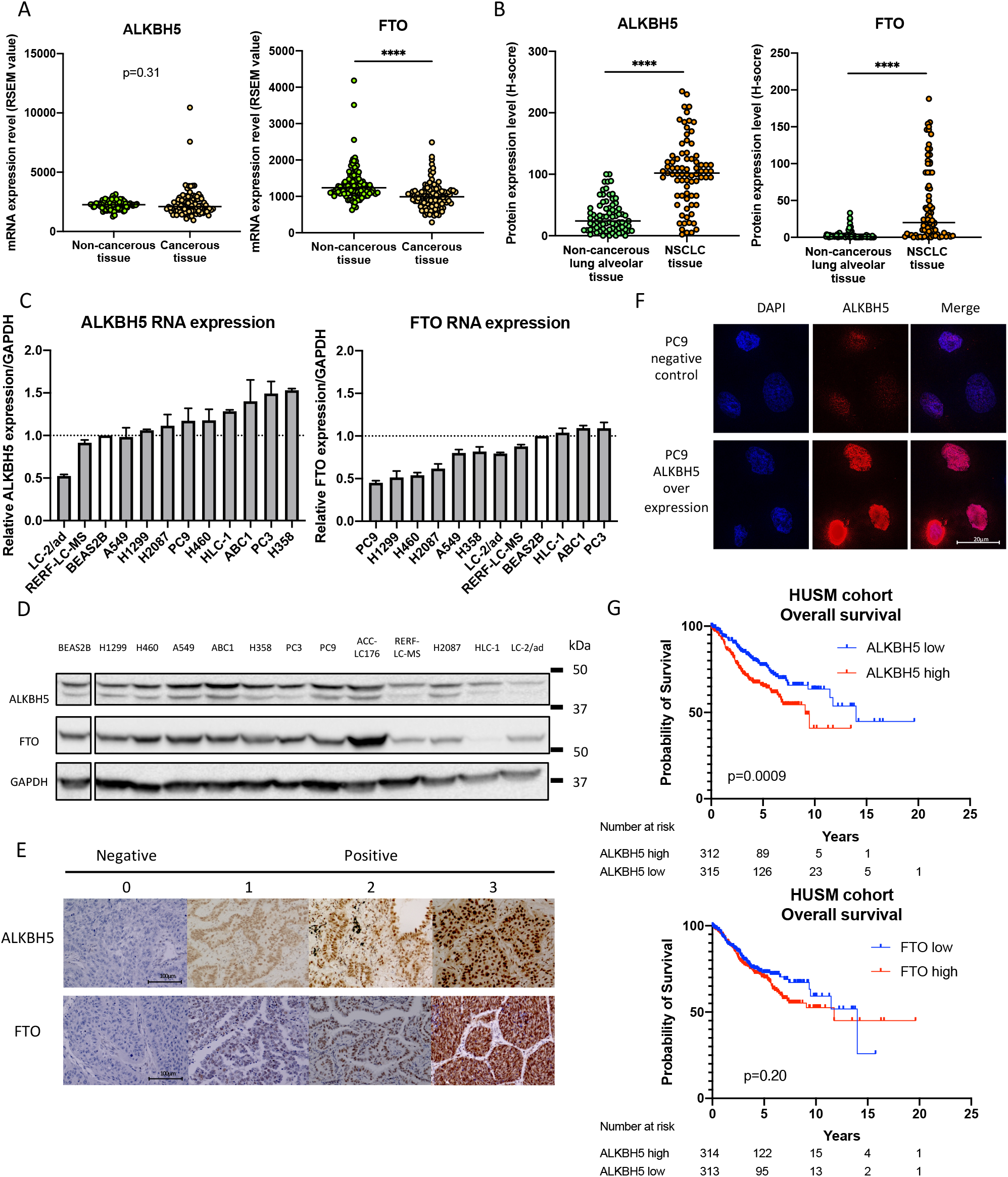
High ALKBH5 expression was associated with a worse prognosis in patients with non-small cell lung cancer. (**a**) ALKBH5 and FTO mRNA levels were analyzed in the paired non-cancerous and NSCLC tissues using the TCGA database (n = 109 for each group). (**b**) ALKBH5 and FTO protein levels were assessed in the paired non-cancerous lung alveolar tissue and NSCLC tissues in the HUSM cohort via immunohistochemistry (IHC) using the H-score (n = 77 for each group). (**c**) Relative ALKBH5 and FTO mRNA expression levels were detected using qPCR in cell lines. Data were normalized to GAPDH and adjusted to the expression of BEAS2B cells (ALKBH5: n = 2, FTO: n = 3). (**d**) ALKBH5 and FTO protein expression levels were determined using western blot analysis in cell lines. (**e**) IHC staining for ALKBH5 and FTO were assessed using the TMA core of NSCLC tissues in the HUSM cohort. Staining intensity was categorized into 0 (absent), 1 (weak), 2 (moderate), or 3 (strong). (**f**) Immunofluorescence visualized subcellular localization in PC9 cells (×100). PC9 cells infected with pRetroX-TetOne puro-ALKBH5 were transduced by 100 ng/mL of doxycycline and used as ALKBH5 overexpression. PC9 cells without doxycycline were used as a negative control. (**g**) A Kaplan–Meier survival curve with log-rank test was utilized to analyze the overall survival of the HUSM cohort. Patients were stratified into low (blue) or high expression groups (red) based on a cutoff determined by the median H-scores (n = 627). Results were presented as the median (a and b) or mean ± SD (c). \*\*\*\**P* < 0.0001 indicates a significant difference between the indicated groups.

### ALKBH5 knockdown suppressed cell proliferation in NSCLC

To investigate the impact of ALKBH5 and FTO deficiency in lung cancer cell function, ALKBH5 and FTO were knocked down in PC9 and A549 cells using small interfering RNA (siRNA). Based on knockdown efficacy, ALKBH5 siRNA no. 1 (siALKBH5#1) and siRNA no. 3 (siALKBH5#3) and FTO siRNA no.1 (siFTO#1) and siRNA no. 3 (siFTO#3) were used in subsequent knockdown experiments (Figure 2a, 2b, and Figure S2A). ALKBH5 knockdown significantly suppressed the proliferation of PC9 and A549 cells (Figure 2c). By contrast, FTO knockdown showed no significant suppressive effects on the proliferation of PC9 and A549 cells (Figure 2d). Thereafter, we assessed migration abilities in ALKBH5-knockdown cells. Accordingly, the transwell migration assay showed no significant reduction in the migratory PC9 and A549 cells (Figure 2e). Moreover, the wound-healing assay showed that ALKBH5 knockdown promoted no significant reduction in the migration ability of PC9 and A549 cells (Figure 2f, 2g). Together with the prognostic value of ALKBH5 in NSCLC, these observations suggested that ALKBH5 played a cancer-promoting role by regulating cell proliferation.

**Figure 2.**
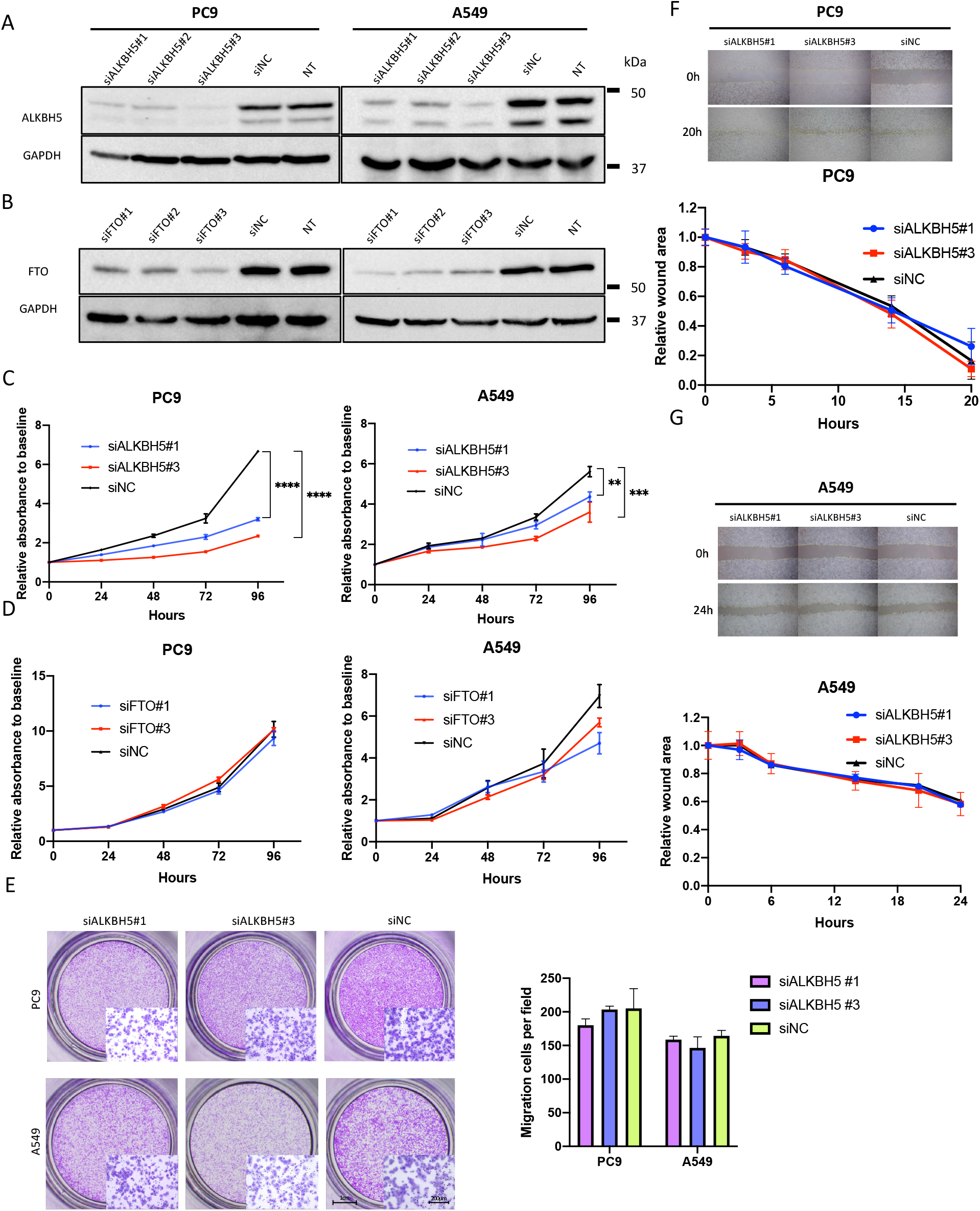
ALKBH5 knockdown suppressed cell proliferation in non-small cell lung cancer. (**a, b**) Western blot analysis demonstrated ALKBH5 protein levels in cells transfected with siRNA for ALKBH5 (siALKBH5), FTO (siFTO), control (siNC), or non-treated cells (NT). (**c**) Cell proliferation relative to baseline in PC9 and A549 cells transfected with siALKBH5 (#1 and #3) or siNC were assessed using the CCK-8 assay (n = 3). **(d)** Cell proliferation relative to baseline in PC9 and A549 cells transfected with siFTO (#1 and #3) or siNC were assessed using the CCK-8 assay (n = 3). (**e**) Migration ability of PC9 and A549 cells transfected with siALKBH5 (#1 and #3) or siNC were assessed using transwell migration assay. The bar charts indicate the number of migratory cells that passed through the chamber membrane (n = 3). (**f, g**) Migration ability of PC9 (**f**) and A549 (**g**) cells transfected with siALKBH5 (#1 and #3) or siNC was assessed using wound-healing assay (n = 3). Results were presented as mean ± SD. \*\**P* < 0.01, \*\*\**P* < 0.001, \*\*\*\**P* < 0.0001 indicates a significant difference between the indicated groups.

To subsequently examine the mechanism by which ALKBH5 knockdown suppressed cell proliferation, cell cycle and apoptosis analyses were performed using flow cytometry. Accordingly, ALKBH5 knockdown significantly increased the number of PC9 cells in the G1 phase and reduced the number of PC9 cells in the G2/M phase (Figure 3a, 3b). Conversely, ALKBH5 knockdown did not reach significant differences, with a consistent result of two different sequences of siRNAs in the cell cycle of A549 (Figure3c, 3d). ALKBH5 knockdown increased the number of apoptotic PC9 cells (Figure 3e, 3f). Furthermore, under drug-induced apoptosis via cisplatin and gefitinib administration, ALKBH5 knockdown also increased the number of apoptotic PC9 cells (Figure 3g, 3h). ALKBH5 knockdown also increased the number of apoptotic A549 cells (Figure 3i, 3j). Moreover, ALKBH5 knockdown increased the number of apoptotic A549 cells with cisplatin (Figure 3k, 3l). Overall, the aforementioned data showed that ALKBH5 knockdown suppressed cell proliferation through G1 phase arrest and/or apoptosis induction in NSCLC cell lines.

**Figure 3.**
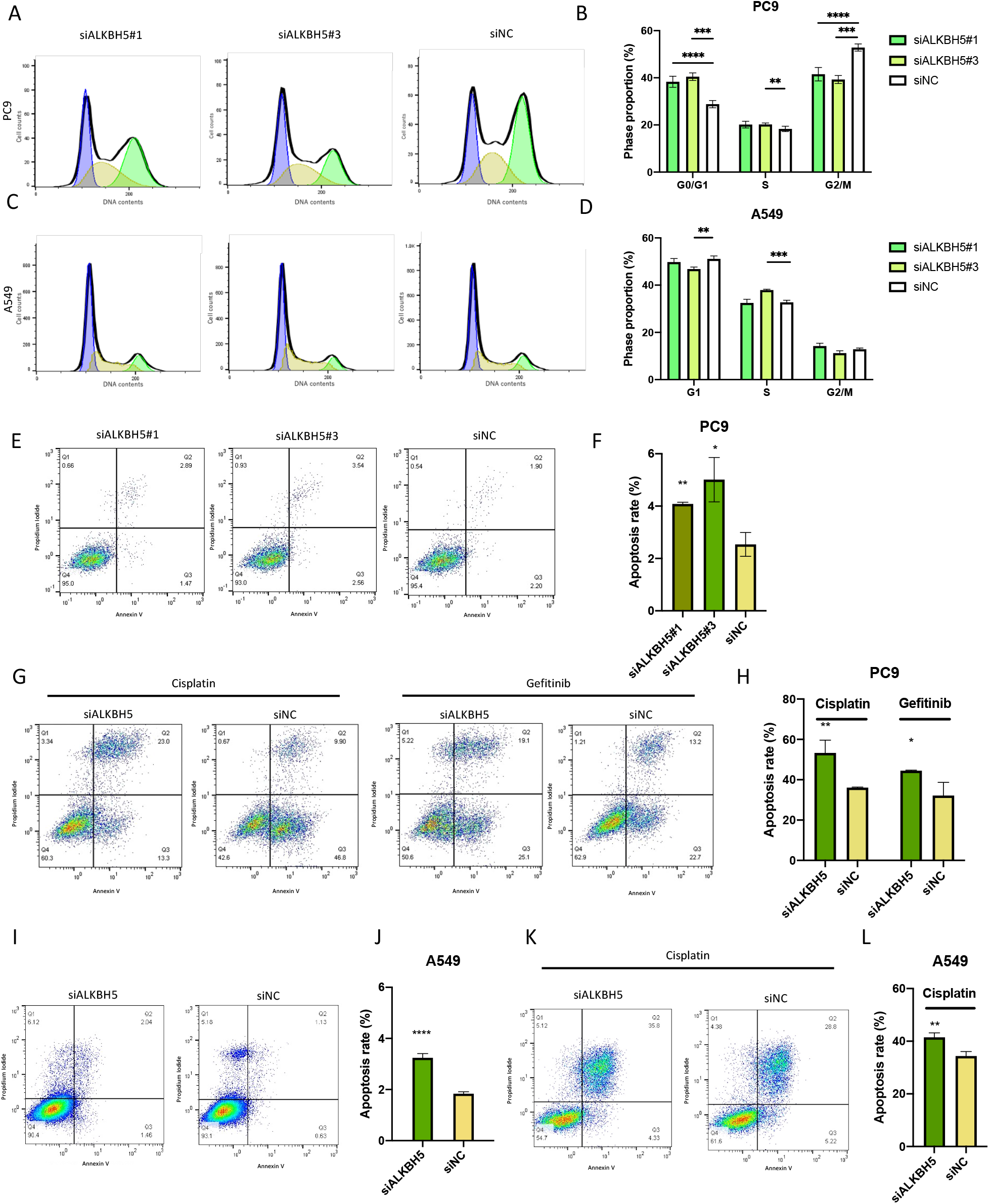
ALKBH5 knockdown-induced G1 phase arrest of cell cycle and/or apoptosis in non-small cell lung cancer. (**a–d**) The cell cycle was examined via flow cytometry with PC9 (a) and A549 (c) cells transfected with siALKBH5 (#1 and #3) or siNC. The bar charts indicate the percentage of cells in each cell cycle phase [n = 6 for PC9 (b), n = 3 for A549 cells (d)]. (**e** and **f**) Apoptotic cells were determined using flow cytometric analysis of PC9 cells transfected with siALKBH5 (#1 and #3) or siNC. (e) Percentage of the apoptotic cells in which both propidium iodide and Annexin V were positive are shown in a representative scatter plot. (f) The apoptosis rate in ALKBH5 knockdown were compared with siNC and shown as a bar chart (n = 3). (**g** and **h**) Cisplatin- or gefitinib-induced apoptotic cells were determined via flow cytometry with PC9 cells transfected with siALKBH5 or siNC. (g) Percentage of apoptotic cells shown in a representative scatter plot. (h) The cisplatin- or gefitinib-induced apoptosis rate in ALKBH5 knockdown were compared with siNC and shown as a bar chart (n = 3). (**i** and **j**) Apoptotic cells were determined via flow cytometry with A549 cells transfected with siALKBH5 or siNC. (i) Percentage of the apoptotic cells shown in a representative scatter plot. (j) Apoptosis rate in ALKBH5 knockdown were compared with siNC and shown as a bar chart (n = 6). (**k** and **l**) Cisplatin-induced apoptotic cells were determined via flow cytometric analysis of A549 cells transfected with siALKBH5 or siNC. (k) Percentage of the apoptotic cells shown in a representative scatter plot. (l) The cisplatin-induced apoptosis rate in ALKBH5 knockdown were compared with siNC and shown as a bar chart (n = 3). Results were presented as mean ± SD. \**P* < 0.05, \*\**P* < 0.01, \*\*\**P* < 0.001, \*\*\*\**P* < 0.0001 indicates a significant difference between the indicated groups.

### ALKBH5 overexpression promoted cell proliferation in immortalized cells

To analyze whether ALKBH5 overexpression in immortalized cells promoted malignant changes in cell function, BEAS2B and HEK293 cells, which are immortalized cells, were infected with a doxycycline-inducible vector, pRetroX-TetOne puro-ALKBH5. ALKBH5 overexpression was confirmed in HEK293 and BEAS2B cells (Figure 4a) and significantly enhanced HEK293 and BEAS2B cell proliferation (Figure 4b and 4c). In contrast, ALKBH5 overexpression showed no significant effects on the migration ability of HEK293 (Figure 4d and 4e) and BEAS2B cells (Figure 4f and 4g). The aforementioned results provided further evidence that ALKBH5 played a cancer-promoting role by regulating cell proliferation.

**Figure 4.**
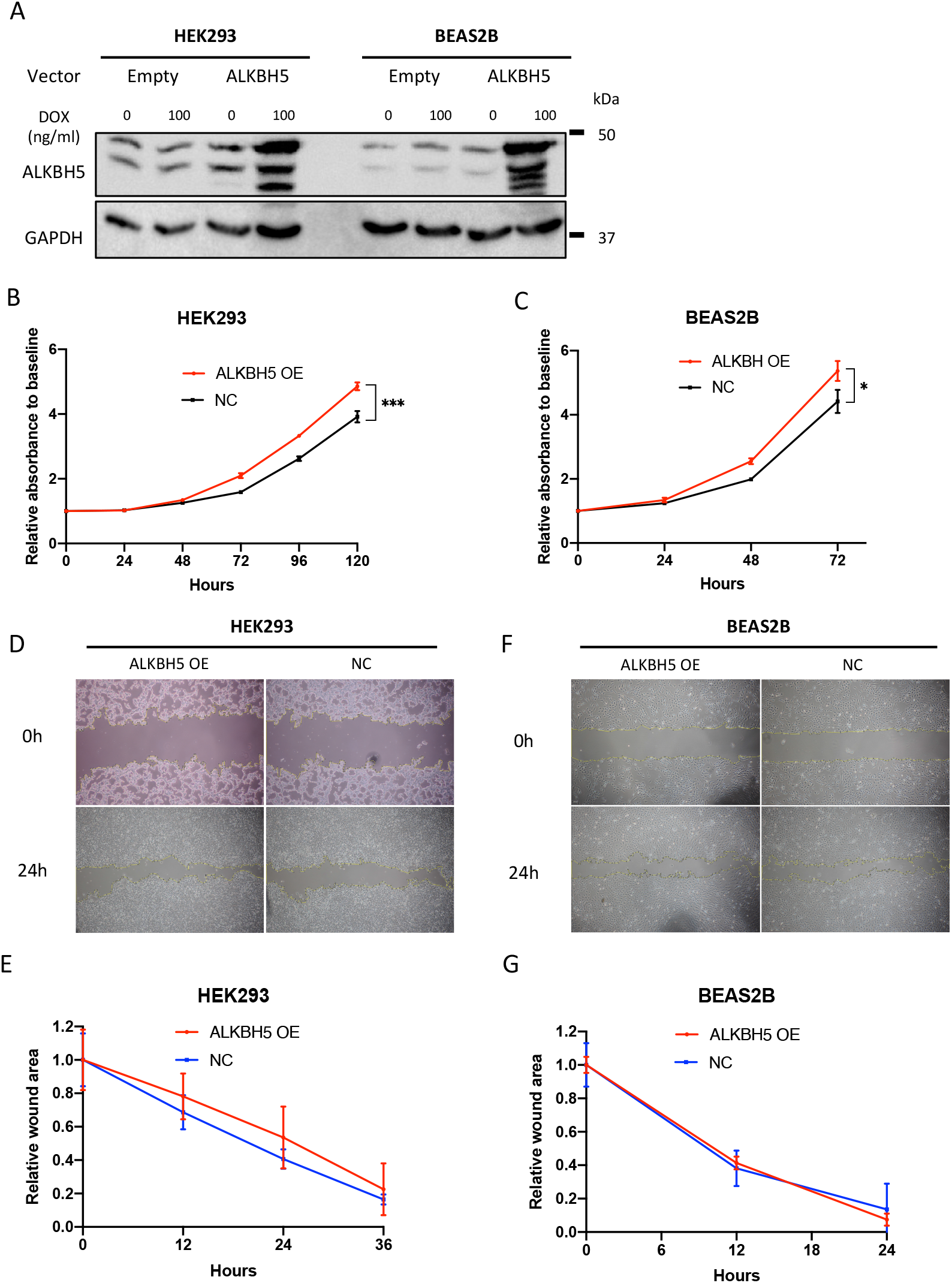
ALKBH5 overexpression promoted cell proliferation. Immortalized cells infected with pRetroX-TetOne puro empty vector (empty) or pRetroX-TetOne puro-ALKBH5 (ALKBH5) were used to assess the function of ALKBH5-overexpressed cells. The concentration of doxycycline (DOX) was 100 ng/mL. (**a**) Western blot analysis demonstrated ALKBH5 protein levels in ALKBH5-overexpressed HEK293 and BEAS2B cells. (**b, c**) Cells infected with pRetroX-TetOne puro-ALKBH5 with DOX were designated as ALKBH5 overexpression (OE), whereas those without DOX were designated as negative control (NC). Cell proliferation relative to baseline in ALKBH5 OE HEK293 and BEAS2B cells were assessed using the CCK-8 assay (n = 3). (**d–g**) The migration ability of HEK293 and BEAS2B cells was assessed via wound-healing assay. Representative images of the wound-healing assay for HEK293 (**d**) and BEAS2B (**f**) cells. Wound areas relative to baseline at each time point were compared between ALKBH5 OE and NC HEK293 (**e**) and BEAS2B (**g**) cells (n = 3). Results are presented as mean ± SD. \**P* < 0.05 indicates a significant difference between the indicated groups.

### ALKBH5 altered the abundance of m^6^A modification in polyA-enriched RNA

To assess the amount of m^6^A in cells, a quantitative evaluation of m^6^A was performed via LC–MS/MS using polyA-enriched RNA extracted from PC9 cells with altered ALKBH5 gene expression (Figure 5a). We investigated the technical variability that occurs when adenosine and *N*^6^-methyladenosine-d3 (m^6^A-d3) are used as internal standards. Although both adenosine (A) and m^6^A-d3 showed a strong positive correlation with m^6^A (r = 0.92 and r = 0.90), the measurement with m^6^A-d3 as the internal standard showed less technical variability than that with A as the internal standard (Figure S3A–C). Hence, we used m^6^A-d3 as the internal control for subsequent experiments. ALKBH5 knockdown increased m^6^A modification in PC9 and A549 cells (Figure 5b and 5c), whereas ALKBH5 overexpression reduced m^6^A modification in a doxycycline concentration-dependent manner in PC9 cells (Figure 5d, S3D, S3E). Moreover, ALKBH5 overexpression reduced m^6^A modification regardless of the time that had elapsed after doxycycline addition (Figure 5e). Furthermore, ALKBH5 overexpression reduced m^6^A modification in BEAS2B and HEK293 cells (Figure 5f). The aforementioned results presented evidence suggesting that ALKBH5 alters the global m^6^A abundance in cells.

**Figure 5.**
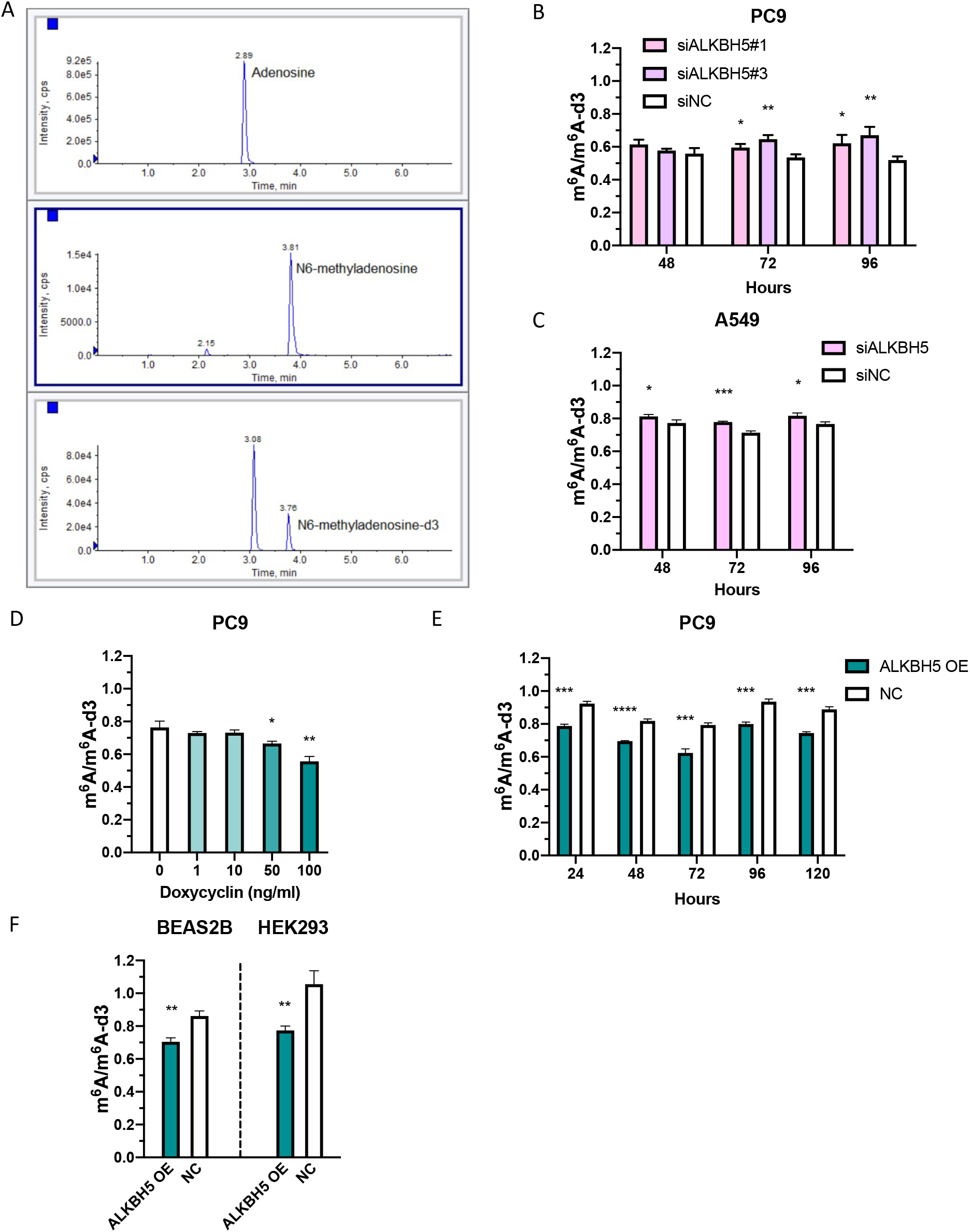
ALKBH5 altered the abundance of m^6^A modification in polyA-enriched RNA. (**a**) Representative chromatograms obtained using liquid chromatography–mass spectrometry/mass spectrometry (LC–MS/MS) for adenosine (upper panel, 2.89 min), *N*^6^-methyladenosine (middle panel, 3.81 min), and *N*^6^-methyladenosine-d3 (lower panel, 3.76 min) in polyA-enriched RNA extracted from PC9 cells. Peak areas were quantified as the product of retention time (min) and count per seconds (cps). (**b** and **c**) The peak areas of m^6^A were normalized to that of m6A-d3. The amount of m^6^A in PC9 (b) and A549 (c) cells transfected with siALKBH5 were compared with that in cells transfected with siNC for 48, 72, and 96 h (n = 3). (**d**) PolyA-enriched RNAs were extracted from PC9 cells containing pRetroX-TetOne puro-ALKBH5 vector. m^6^A/m^6^A-d3 in the cells incubated with various concentrations of DOX for 48 h were compared with that in cells incubated with 0 ng/mL DOX (n = 3). (**e**) m^6^A/m^6^A-d3 in PC9 cells containing pRetroX-TetOne puro-ALKBH5 vector (ALKBH5 OE) whose ALKBH5 overexpression was induced by 100 ng/mL DOX for 24, 48, 72, 96, 120 h was compared with those without DOX (NC) (n = 3). (**f**) m^6^A/m^6^A-d3 in HEK293 and BEAS2B (ALKBH5 OE) cells whose ALKBH5 overexpression was induced by 100 ng/mL DOX for 48 h were compared with those without DOX (NC) (n = 3). Results were presented as mean ± SD. \**P* < 0.05, \*\**P* < 0.01, \*\*\**P* < 0.001, \*\*\*\**P* < 0.0001 indicates a significant difference between the indicated groups by Student’s t-test.

### ALKBH5 regulated the expression of cell proliferation-related genes

An expression microarray analysis was herein performed to investigate gene expression profiles in ALKBH5-knockdown PC9 cells with two different sequences of siRNA (ALKBH5#1 and ALKBH5#3). Differentially expressed genes (DEGs) were defined as those with a fold change of >1.5 or <0.67 (*P* < 0.01). A total of 697 DEGs were detected for ALKBH5#1 comprising 392 upregulated and 305 downregulated genes (Figure 6a), whereas 1394 DEGs were detected for ALKBH5#3 comprising 803 upregulated and 591 downregulated genes (Figure 6b). Moreover, 82 upregulated genes (Table S4) and 47 downregulated genes (Table S5) overlapped between ALKBH5#1 and ALKBH5#3 (Figure 6c). Except ALKBH5, genes associated with m^6^A modification described in a previous review [47] were not included in the overlapped DEGs (Table S6). GSEA with the hallmark gene set revealed that the PC9 cells transfected with siALKBH5#1 and siALKBH5#3 had a more enriched expression of genes involved in cell cycle, such as MYC_TARGETS_V2, P53_PATHWAY, and G2/M_CHECKPOINT, than those transfected with siNC (Figure S4A and S4B). We selected 10 DEGs associated with cell proliferation bibliographically and confirmed the upregulation of E2F1, GADD45A, TIMP3, and CDKN1A and downregulation of CASP14 and CCNG2 by qPCR (Figure 6d). The aforementioned results revealed that ALKBH5 regulated the expression of genes associated with cell proliferation.

**Figure 6.**
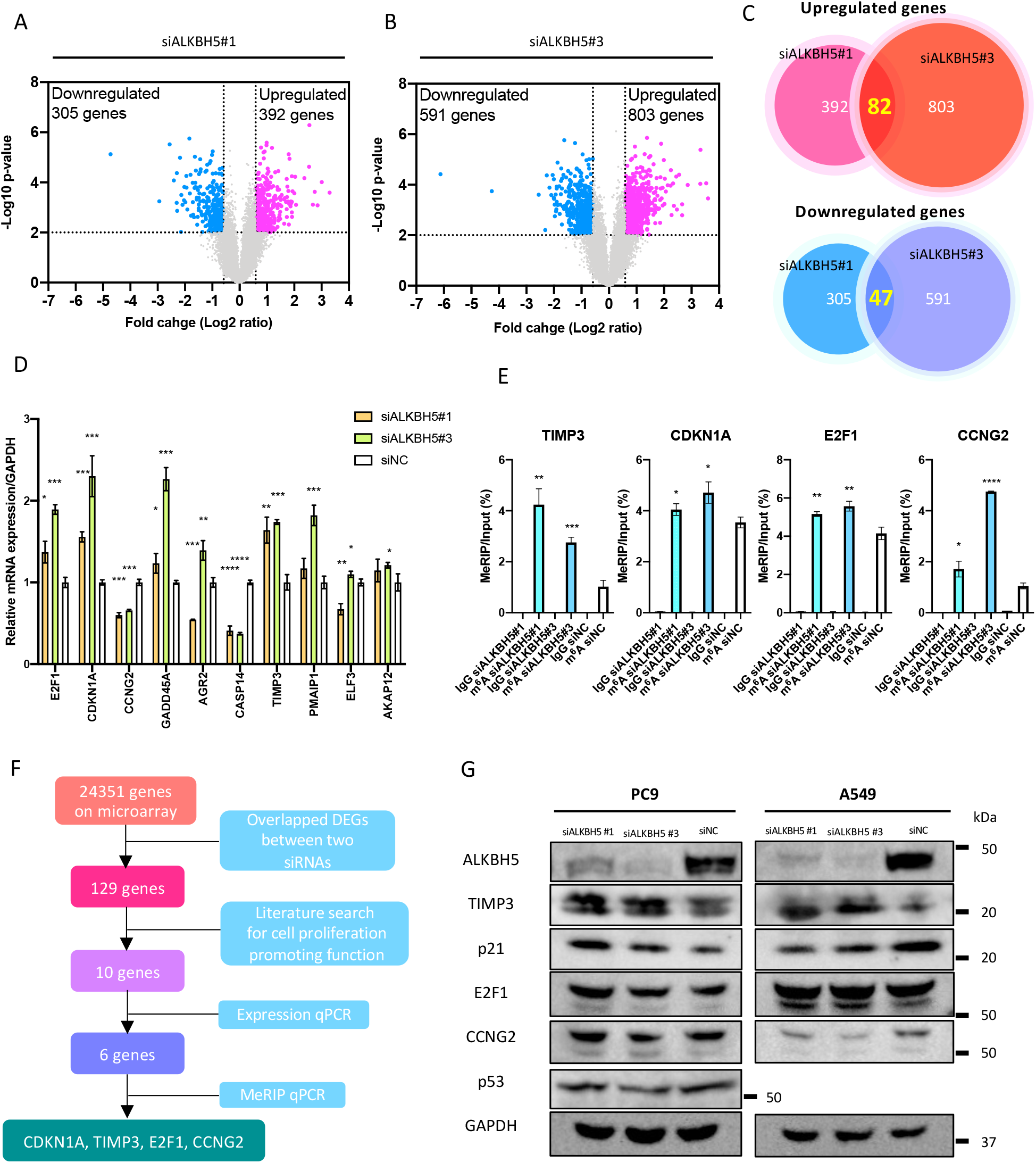
ALKBH5 knockdown regulated cell proliferation-related genes and m^6^A abundance in the 3′ untranslated regions of specific genes. PC9 cells were transfected with siNC, siALKBH5#1, or siALKBH5#3 for 96 h (n = 3 for each group). (**a, b**) Differentially expressed genes (DEGs) for siALKBH5#1 (a) or siALKBH5#3 (b) were detected using expression microarray and shown using volcano plots. Dashed lines indicate the threshold for the differential expression [fold change > 1.5 (log2 fold change = 0.5849) or < 0.67 (log2 fold change = −0.5849), *P* < 0.01 via Student’s t-test] for upregulated (pink dots) or downregulated (light blue dots) genes. (**c**) Venn diagram indicating the number of common DEGs in ALKBH5-knockdown cells with different siRNA sequences. (**d**) mRNA expression levels of genes related to cell proliferation in ALKBH5-knockdown PC9 cells were analyzed using qPCR. Gene expression was normalized to the GAPDH expression and was shown relative to the expression with siNC. (**e**) m^6^A level in the 3′ UTRs of target mRNA in PC9 cells transfected with siALKBH5#1 or siALKBH5#3 was quantified via MeRIP qPCR using anti-m6A antibody and was compared with that in cell transfected with siNC. The m^6^A level was normalized to that of input fraction (n = 3). IgG was used to evaluate the non-specific binding of the target mRNA. (**f**) A schematic outline showing the workflow for the analysis of downstream targets of ALKBH5. (**g**) Target protein levels in PC9 or A549 cells transfected with siALKBH5#1 or siALKBH5#3 were compared with those transfected with siNC via western blot analysis. Results are presented as mean ± SD. \**P* < 0.05, \*\**P* < 0.01, \*\*\**P* < 0.001, \*\*\*\**P* < 0.0001 indicates a significant difference between the indicated groups.

### ALKBH5 altered the abundance of m6A in the 3**′** UTR and regulated protein expression of target genes

We performed m^6^A-specific methylated RNA immunoprecipitation microarray analysis in PC9 cells on an Arraystar Human mRNA&lncRNA Epitranscriptomic Microarray to comprehensively examine whether differentially regulated genes were associated with m^6^A modification using unfragmented total RNA. The median methylation level in unfragmented total RNA was 50.4% (6.8%–94.5%) (Figure S5A). A positive correlation was observed between the methylation level in unfragmented total RNA and the RNA length of each transcript (r = 0.35) (Figure S5B). Moreover, a negative correlation was noted between the rate at which ALKBH5 knockdown increased m^6^A modification (methylation level in siALKBH5 − methylation level in siNC) and methylation level at baseline (methylation level in siNC) (r = −0.35) (Figure S5C). The volcano plot showed that 1 RNA was hypermethylated by ALKBH5#1 knockdown (fold change > 1.5, *P* < 0.01) (Figure S5D), whereas 28 RNAs were hypermethylated by ALKBH5#3 knockdown (fold change > 1.5; *P* < 0.01) (Figure S5E). No hypermethylated genes overlapped between ALKBH5#1 and ALKBH5#3 knockdown with a fold change threshold of >1.5 (*P* < 0.01) (Figure S5F). GSEA showed no common hallmark gene set with an FDR q-value of < 0.25 for PC9 cells transfected with siALKBH5#1 and siALKBH5#3 compared with the control group (Figure S5G).

Considering that the m6A levels of unfragmented RNAs regulated by ALKBH5 is affected by the baseline RNA length and endogenous m6A level, we performed methylated RNA immunoprecipitation (MeRIP) with m^6^A antibody using fragmented polyA-enriched RNA in ALKBH5-knockdown PC9 cells to investigate focal m^6^A alterations in the mRNA. The fragmentation conditions were optimized (Figure S6A), and the m^6^A changes in the positive and NC were confirmed through qPCR using the primers included in the Magna MeRIP m^6^A Kit (Millipore) (Figure S6B). To verify the accuracy of the MeRIP experiment, we selected MFAP5 out of the 11 genes that were hypermethylated (>1.5 fold change and *P* < 0.05) in the human mRNA&lncRNA Epitranscriptomic Microarray (Table S7) and analyzed the m^6^A target site via qPCR. Specific primers were designed for the predicted m^6^A-harboring regions, and MeRIP qPCR confirmed that ALKBH5 knockdown increased m^6^A levels in the 3′ UTR of MFAP5 (Figure S6C). Thereafter, MeRIP qPCR was performed in six DEGs verified using qPCR with ALKBH5-knockdown PC9 cells. Our results showed increased m^6^A levels in the 3′ UTRs of CDKN1A, TIMP3, E2F1, and CCNG2 in ALKBH5-knockdown PC9 cells (Figure 6e and Figure S6D). The aforementioned results of the MeRIP qPCR suggest that ALKBH5 targeted the 3′ UTRs of m A in these four transcripts (Figure 6f).

Next, the protein expression levels of the potential target transcript of ALKBH5 were quantified by western blot analysis. Accordingly, our results showed that CDKN1A (p21) expression increased independent of p53 in ALKBH5-knockdown PC9 cells, whereas TIMP3 expression increased in ALKBH5-knockdown A549 cells (Figure 6G).

### IGF2BPs were required for ALKBH5 regulation of target mRNA expression via stabilization of mRNA and affected cell proliferation

IGF2BP1, IGF2BP2, and IGF2BP3 (IGF2BPs) are well-known m^6^A-recognizing RNA-binding proteins and readers of m^6^A that have been known to stabilize mRNA. The expression of these proteins in a series of cell lines was analyzed by western blotting, and the results showed significant differences in their expression in lung cancer cells and immortalized bronchial epithelial cells according to the cell lines (Figure 7a).

**Figure 7.**
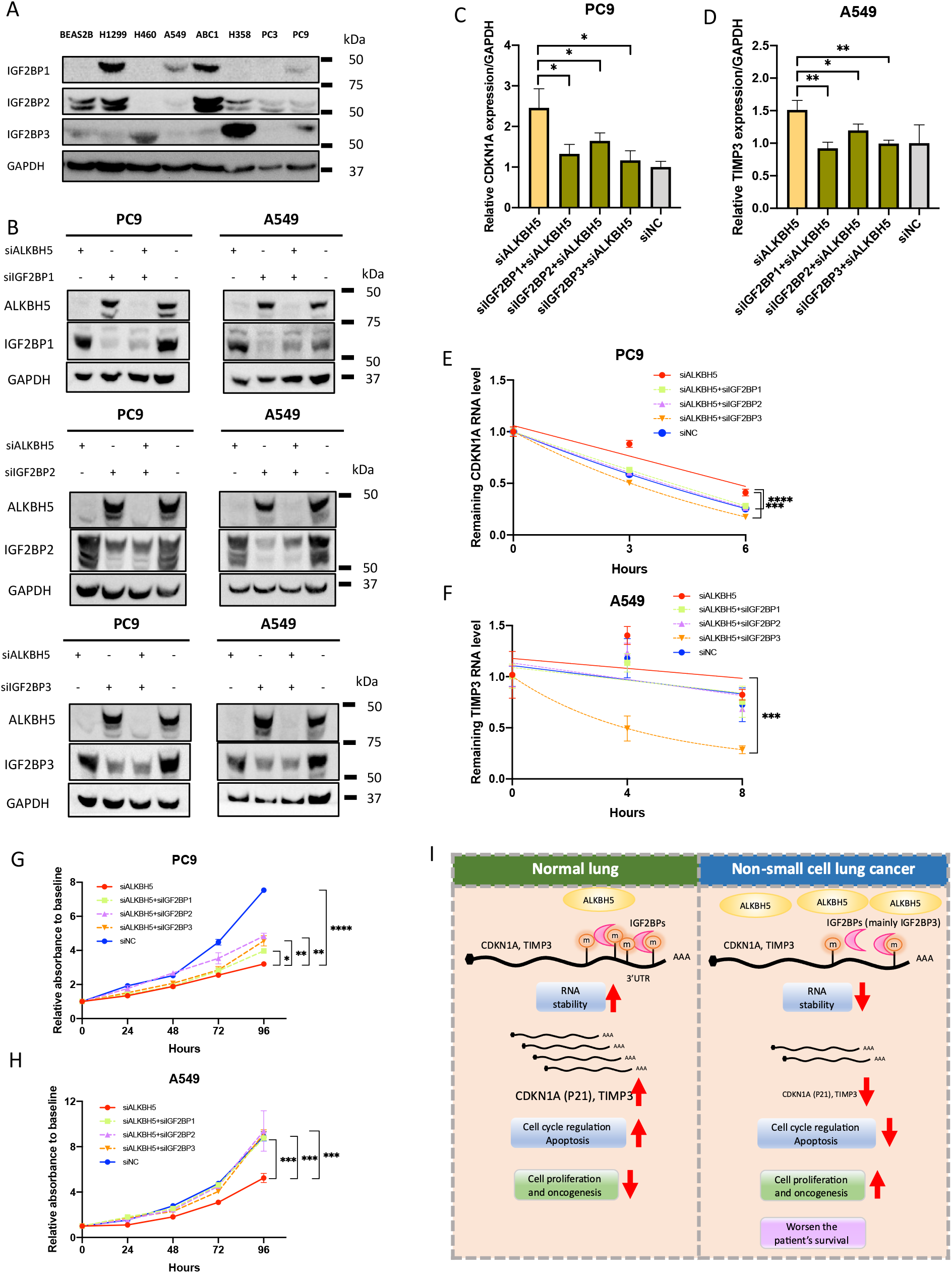
IGF2BPs was required for the ALKBH5-induced regulation of mRNA expression and cell proliferation. (**a**) Expression levels of IGF2BP1, IGF2BP2, and IGF2BP3 (IGF2BPs) protein determined using western blot analysis were compared between cell lines. (**b**) IGF2BP protein levels in cells transfected with siALKBH5 (left end), those in cells transfected with siIGF2BP1, siIGF2BP2, or siIGF2BP3 with or without siALKBH5 (middle two lanes), or those with siNC (right end indicating both siALKBH5 and siIGF2BPs were negative) were confirmed via western blot analysis. (**c** and **d**) Endogenous mRNA expression levels of CDKN1A in PC9 cells (c) or those of TIMP3 in A549 cells (d) transfected with siALKBH5 were analyzed via qPCR and compared with those in cells co-transfected with siALKBH5 and one of the siIGF2BPs. Gene expression was normalized to the GAPDH expression and was shown relative to the expression in siNC (n = 3). (**e** and **f**) The remaining RNA level of CDKN1A in PC9 cells (e) or of TIMP3 in A549 cells (f) after actinomysin D treatment for 0, 2, 4, and 6 h was determined using qPCR and normalized to the expression at 0 h. RNA decay rate in cells transfected with siALKBH5 and/or one of the siIGF2BPs and siNC were compared with the stability of CDKN1A and TIMPs (n = 3). (**g** and **h**) Cell proliferation relative to baseline in PC9 (g) and A549 (h) cells transfected with siALKBH5 was assessed via the CCK-8 assay and compared with that in cells cotransfected with siALKBH5 and one of the siIGF2BPs (n = 3 for each group). (**i**) Schematic illustration for the proposed mechanism of tumorigenicity via ALKBH5 in non-small cell lung cancer. Upregulation of ALKBH5 in NSCLC reduces m^6^A modifications on the 3′ UTR of specific genes. The loss of m^6^A decreases the opportunity for recognition by IGF2BPs and destabilizes the target transcript, such as CDKN1A (p21) and TIMP3. Downregulation of CDKN1A (p21) and TIMP3 induces cell cycle alteration and inhibits apoptosis. This ALKBH5–IGF2BPs axis promotes cell proliferation and tumorigenicity, which in turn causes the unfavorable prognosis of NSCLC. Results are presented as mean ± SD. \**P* < 0.05, \*\**P* < 0.01, \*\*\**P* < 0.001, \*\*\*\**P* < 0.0001 indicates a significant difference between the indicated groups.

Thereafter, ALKBH5 and IGF2BPs were knocked down with siRNA to investigate the association between ALKBH5 and IGF2BPs and the expression of CDKN1A or TIMP3 (Figure S7A, S7B, S7C, S7D, 7b). Accordingly, ALKBH5 knockdown increased the mRNA expressions of CDKN1A in PC9 cells (Figure 7c) and TIMP3 in A549 cells (Figure 7d). In addition, the increased expression was offset by the knockdown of IGF2BPs (Figure 7c and 7d). Actinomycin D assay showed that ALKBH5 knockdown stabilized CDKN1A mRNA in PC9 cells, and this stabilization was offset by the knockdown of IGF2BPs (Figure 7e). ALKBH5 knockdown also stabilized TIMP3 mRNA in A549 cells, although not statistically significant, and this stabilization was decreased by IGF2BP3 knockdown (Figure 7f). These results suggest that these alterations in mRNA expression were offset by a double knockdown of both ALKBH5 and one of the IGF2BPs, and the decline of mRNAs were, at least partly, owing to the destabilization of these mRNAs by one of the IGF2BPs. Additionally, we evaluated the expression of ALKBH5 target mRNAs CDKN1A and TIMP3 in lung cancer using the TCGA dataset. Accordingly, cancerous tissues (high ALKBH5 expression) had lower CDKN1A and TIMP3 expression than non-cancerous tissue (low ALKBH5 expression) (Figure S7E and S7F).

Considering that the interaction between ALKBH5 and IGF2BPs was found to regulate the expression of genes associated with cell proliferation, cell proliferation assays were conducted using ALKBH5- and IGF2BPs-knockdown cells. Notably, ALKBH5 knockdown reduced cell proliferation in PC9 (Figure 7g) and A549 cells (Figure 7h). The reduction of cell proliferation was offset by IGF2BPs knockdown. These results support the hypothesis that IGF2BPs are required for ALKBH5 regulation of target mRNA expression and cell proliferation.

## Discussion

The current study revealed that ALKBH5 promoted poor survival and cell proliferation in patients with NSCLC. Mechanistically, ALKBH5 knockdown had been found to increase the expression of CDKN1A (p21) and TIMP3 by altering mRNA stability in PC9 and A549 cells via m^6^A change. Moreover, these changes in mRNA stability were counteracted by IGF2BPs knockdown (particularly prominent in IGF2BP3). The aforementioned results suggest that the recognition of target transcripts by IGF2BPs stabilizes the mRNA of CDKN1A (p21) or TIMP3 and subsequently increases their expressions, thereby regulating cell proliferation, cell cycle, and apoptosis in lung cancer cell lines.

Over the last decade, considerable progression has been made on research regarding the molecular mechanism for m^6^A-mediated carcinogenesis of ALKBH5. Nevertheless, previous studies on ALKBH5 have shown conflicting results regarding the carcinogenic mechanisms of ALKBH5 across several cancers [28–36]. Two previous studies have reported contradictory results regarding ALKBH5, suggesting that it acts as either an oncogenic factor or a tumor suppressor in NSCLC [40, 41]. The current study concluded that ALKBH5 exerted cancer-promoting effects in NSCLC by suppressing CDKN1A (p21) or TIMP3. CDKN1A (p21) functions as a cell growth suppressor by inhibiting cell cycle progression. Multiple transcription factors, ubiquitin ligases, and protein kinases regulate the transcription, stability, and cellular localization of CDKN1A (p21) [48]. A previous study showed that ALKBH5 knockdown increased m^6^A modification and mRNA stability of CDKN1A, which subsequently increased p21 protein expression and acted as a tumor suppressor in esophageal cancer [32]. Similarly, our findings showed that ALKBH5 knockdown in PC9 cells acted as a tumor suppressor by the upregulation of CDKN1A (p21) via m^6^A alteration. We also showed that p21 expression was p53-independent, indirectly reinforcing our finding that CDKN1A (p21) upregulation was an m^6^A-mediated response. Our results further indicated a novel mechanism wherein changes in CDKN1A expression via ALKBH5 knockdown were rescued by IGF2BPs knockdown, which supports our finding that alterations in CDKN1A (p21) expression were mediated by m^6^A.

The current study identified TIMP3 as another important target molecule downstream of ALKBH5. A previous study showed TIMP3 had several anticancer properties, including apoptosis induction and antiproliferative, antiangiogenic, and antimetastatic activities. The expression of TIMP3 is regulated by transcription factors and histone acetylation [49]. Several studies have shown that TIMP3 acts as a tumor suppressor in lung cancer [50, 51]. Moreover, a previous report using A549 cell lines showed that ALKBH5 knockdown increases TIMP3 mRNA stability and TIMP3 expression via m^6^A modification [41]. Similarly, the current study also confirmed that ALKBH5 knockdown increased mRNA stability, which increased TIMP3 protein expression, and acted as a tumor suppressor in A549 cells. Furthermore, our experimental data for the first time showed that the ALKBH5 knockdown-induced increase in TIMP3 was rescued by IGF2BPs, strongly suggesting that alterations in TIMP3 expression were mediated by m^6^A.

Previous studies have reported that IGF2BPs, which are known as m^6^A-recognizing RNA-binding proteins that stabilize m^6^A-containing RNA, have oncogenic properties. Studies on lung cancer have associated IGF2BPs with cancer progression and poor prognosis [52–57]. Notably, a previous report showed that ALKBH5-mediated m^6^A modification of LY6/PLAUR Domain Containing 1 (LYPD1) is recognized by IGF2BP1 and enhances the stability of LYPD1 mRNA in hepatocellular carcinoma [35]. Moreover, recent RNA-binding protein immunoprecipitation-sequencing analysis using HEK293T showed that the binding site of IGF2BPs is mainly distributed in the 3′ UTRs and that the target of IGF2BPs preferentially binds to the consensus sequence of UGGAC in the target mRNA [26]. These findings support our experimental hypothesis that IGF2BPs recognize the m^6^A in the 3′ UTRs of CDKN1A or TIMP3 because UGGAC is present within three locations in the 3′ UTRs of CDKN1A and two locations in the 3′ UTRs of TIMP3. Our findings also showed that the expression of IGF2BPs significantly differed with the lung cancer cell line, which can cause different fates of m^6^A-modified transcripts among cell lines.

FTO inhibitors have been shown to suppress the progression of acute myeloid leukemia and glioblastoma *in vivo* [58, 59]. In contrast, the antitumor effects of ALKBH5 inhibitors, which enhanced the efficacy of cancer immunotherapy, have only been confirmed in melanomas [60]. Furthermore, several experimental facts have shown that ALKBH5 is associated with the malignant transformation of cancer [28–34], indicating that ALKBH5 inhibitors can be a target of tumor-agnostic therapy. However, it should be noted that ALKBH5 inhibitors may cause unexpected side effects in unknown target genes given that ALKBH5 inhibition alters the m6A modification of numerous transcripts and the expression of several genes.

Although the current study provided abundant evidence to conclude the remarkable role of the m^6^A-regulated ALKBH5 and IGF2BPs axis in NSCLC, several limitations warrant consideration. First, given that RNAs were not fragmented during our epitranscriptomic microarray, site-specific changes in m^6^A modification could not be determined. Considering the correlation between RNA length and ratio of m^6^A-modified transcripts, the epitranscriptional micrarray analysis using unfragmented RNA does not allow the evaluation of multiple m6A modifications occurring within a single transcript. As such, we conducted MeRIP q-PCR with fragmented RNAs to evaluate site-specific differential m^6^A modification. Nevertheless, the epitranscriptomic microarray with unfragmented RNAs provided a holistic view of the degree of m^6^A modification for each transcript, establishing a landscape for m^6^A modification by ALKBH5 knockdown (Figure S5A, S5B, S5C, S5D, S5E, and S5G). Secondly, our epitranscriptomic microarray findings showed that ALKBH5-knockdown reduced m6A methylation levels in approximately half of the transcripts. Although the detailed mechanism remains unclear, hypomethylation may occur when some of the m^6^A-rich transcripts bind to YTHDF2 and YTHDC2, reducing the stability of RNA containing m^6^A. Consequently, the m^6^A-modified transcript then undergoes degradation. In other words, the target’s transcript may also differ depending on the elapsed time after the perturbation of ALKBH5. However, the current study did not investigate the chronological alteration of the m^6^A abundance of each transcript following ALKBH5 knockdown. Third, as mentioned earlier, CDKN1A and TIMP3 are also regulated by transcription factors or miRNA, and we cannot deny the possibility that mechanisms other than m^6^A promoted changes in CDKN1A (p21) and TIMP3 expression. Nonetheless, the finding that IGF2BPs knockdown rescued the CDKN1A and TIMP3 expression supports our proposition that the changes in CDKN1A (p21) and TIMP3 expression were mediated by m^6^A.

## Conclusions

The current study revealed that increased ALKBH5 expression was an independent unfavorable prognostic factor in NSCLC. Moreover, upregulation of ALKBH5 in NSCLC reduced m^6^A modifications on the 3′ UTR of specific genes. The loss of m^6^A decreased the opportunity for recognition by IGF2BPs and destabilized the target transcript, such as CDKN1A (p21) and TIMP3. Downregulation of CDKN1A (p21) and TIMP3 induced cell cycle alteration and inhibited apoptosis. Our results suggest that the ALKBH5–IGF2BPs axis promotes cell proliferation and tumorigenicity, which in turn causes the unfavorable prognosis of NSCLC. Our findings provide a novel insight into the pathophysiological mechanisms of m^6^A epitranscriptomic modification in NSCLC (Figure 7i). Further *in vivo* studies are nonetheless required to determine whether ALKBH5 inhibitors can be incorporated in the treatment of NSCLC in the near future.

## List of abbreviations

m^6^A: N^6^-methyladenosine
NSCLC: non-small cell lung cancer
MeRIP: Methylated RNA immunoprecipitation
CI: Confidence interval
HR: Hazard ratio
UTRs: 3’ untranslated regions
METTL3: methyltransferase-like 3
METTL14: Methyltransferase-like 14
WTAP: Wilms tumor 1-associated protein
RBM15: RNA-binding motif protein 15
FTO: fat mass and obesity-related protein
ALKBH5: AlkB homolog 5
YTHDC: YT521-B homology domain containing
HNRNPG: heterogeneous nuclear ribonucleoprotein G
YTHDF: YT521-B homology domain family
eIF3: eukaryotic initiation factor3
IGF2BP: insulin-like growth factor 2 mRNA-binding protein
m^6^Am: N^6^,2’ -O-dimethyladenosine
TMA: Tissue microarray
IHC: immunohistochemistry
TCGA: the Cancer Genome Atlas
RSEM: RNA-seq by Expectation Maximization
DAPI: 4’,6-diamidino-2-phenylindole
siNC: siRNA control
NT: non-treated cells
DOX: doxycycline
OE: overexpression
NC: negative control
CCK-8: Cell Counting Kit-8
SDS: sodium dodecyl sulfate
PI: propidium iodide
LC–MS/MS: liquid chromatography–mass spectrometry/mass spectrometry
m^6^A-d3: N6-methyladenosine-d3
A: adenosine
FDR: false discovery rate
GSEA: Gene set enrichment analysis
GEO: Gene Expression Omnibus
NCBI: National Center for Biotechnology Information
SD: standard deviation
OS: Overall survival
RFS: recurrence-free survival
ANOVA: analysis of variance
DEGs: differentially expressed genes
LYPD1: LY6/PLAUR Domain Containing 1

## Declarations

### Acknowledgments

We are grateful to the patients and sample donors for their dedicated participation in this study. We also thank Takaharu Kamo, Shiho Omori, Hisaki Igarashi (Tumor Pathology, Hamamatsu University School of Medicine), Ryo Horiguchi, Masako Suzuki, and Takuya Kitamoto (Advanced Research Facilities and Services, Hamamatsu University School of Medicine) for providing technical assistance.

### Author contributions

HS conceived this project, supervised all experiments and interpretations, and drafted the manuscript. KT designed and performed all experiments, analyzed the data, interpreted patient and experimental data, and drafted the manuscript. KY designed this project, performed sample collection, histological examination, and a part of the experiments and drafted the manuscript. Yusuke Inoue performed sample collection and histological examination. Yuji Iwashita designed this project, interpreted the experimental data, and drafted the manuscript. TO performed molecular experiments, interpreted the experimental data, and drafted the manuscript. HY and HW assisted part of the experiments. AK, MT, HO, and KF performed sample collection. KS performed sample collection and analyzed data. TS supervised this project. All authors have read and approved the final manuscript.

### Funding

This work was supported by grants from the Japan Society for the Promotion of Science (JP22659072, JP24659161, JP26670187, JP16K15256) and HUSM Grant-in-Aid, and Smoking Research Foundation.

### Availability of data and materials

All data and supplementary information within the article are available from the published article (including supplementary information files) or available on published databases (TCGA or GEO). GEO accession numbers of our microarray data are GSE165453 and GSE165453.

### Ethics approval and consent to participate

This study was approved by the Ethics Committees of Hamamatsu University School of Medicine (20-011) and Seirei Mikatahara General Hospital and was carried out in accordance with approved guidelines. Written informed consent was obtained from all patients. All analyses were conducted in compliance with the ethical standards according to the Helsinki Declaration.

### Consent for publication

Not applicable.

### Competing interests

The authors declare no competing interests.

## Notes

### Competing Interest Statement

The authors have declared no competing interest.

