## Supplementary text for "m^6^A demethylase ALKBH5 promotes tumor cell proliferation by destabilizing IGF2BPs target genes and worsens the prognosis of patients with non-small cell lung cancer"

**Additional file 1: Figure S1. High ALKBH5 expression was associated with a worse prognosis in patients with NSCLC (related to Figure 1).**

(**a**) ALKBH5 and FTO protein expression level in NSCLC of the HUSM cohort (n = 627) were quantified via immunohistochemistry using the H-score. Correlations were evaluated using the Spearman’s correlation coefficient (r = 0.41). (**b** and **c**) Kaplan–Meier survival curves with log-rank test for ALKBH5 (b) and FTO (c) were applied for prognostic evaluation of recurrence-free survival in the HUSM cohort (n = 627). Patients were stratified into low (blue) or high (red) expression groups based on a cutoff determined by the median H-scores. (**d** and **e**) Kaplan–Meier survival curves with log-rank test for ALKBH5 (d: n = 1144) and FTO (e: n = 1925) were applied for prognostic evaluation of overall survival using the Kaplan–Meier plotter. Patients were stratified based on the median expression of each mRNA.

**Additional file 2: Figure S2. Knockdown efficacy of siRNAs for ALKBH5 and FTO (related to Figure 2).**

(**a**) Relative ALKBH5 and FTO mRNA expression levels were detected via qPCR in PC9 and A549 cells transfected with siALKBH5, siFTO, or siNC. Gene expression was normalized to the expression of GAPDH and shown relative to siNC (n = 1).

**Additional file 3: Figure S3. Technical variability in m^6^A measurement using LC–MS/MS and confirmation of ALKBH5 protein induction (related to Figure 5).**
(**a**) Chemical structural formula of *N*^6^-methyladenosine-d3 (m^6^A-d3). (**b**) Amount of m^6^A in polyA-enriched RNAs in various cell lines (n = 27 samples) was quantified using LC–MS/MS and normalized as m^6^A/m^6^A-d3 or m^6^A/A. Coefficients of variation between the technical duplicate of each sample were plotted and compared using two different methods of normalization. (**c**) Correlations between m^6^A and m^6^A-d3 (left panel, r = 0.92) and between m^6^A and adenosine (right panel, r = 0.90) in polyA-enriched RNAs (n = 54) were evaluated using Spearman’s correlation coefficient. (**d**) PC9 cells were infected with pRetroX-TetOne puro vector (empty) or pRetroX-TetOne puro-ALKBH5 (ALKBH5) and incubated with 0 or 100 ng/mL of DOX for 48 h. ALKBH5 protein levels were quantified via western blot analysis. Cells without the infection process were used as the vector (−). (**e**) PC9 cells infected with pRetroX-TetOne puro-ALKBH5 were incubated with 0–100 ng/mL of DOX for 48 h. ALKBH5 protein levels were quantified via western blot analysis. Bars indicate the median. *****P* < 0.0001 indicates a significant difference between the indicated groups.

**Additional file 4: Figure S4. Gene set enrichment analysis showed cell proliferation related gene sets (related to Figure 6).**

(**a**) Expression profiles of PC9 cells transfected with siALKBH5#1 (in the left panel) and siALKBH5#3 (in the right panel) were compared with those transfected with siNC via gene set enrichment analysis using hallmark gene sets. Significantly enriched gene sets were defined as those with FDR < 0.25. Common enrichment gene sets between PC9 cells transfected with siALKBH5#1 and siALKBH5#3 were presented in the right panel. (**b**) The ranks of genes involved in MYC_TARGETS_V2, P53_PATHWAY and G2/M_CHECKPOINT (black vertical bars) are enriched in highly upregulated genes through both ALKBH5#1(left side) and ALKBH5 #3 (right side) knockdown (red side).

**Additional file 5: Figure S5. Comprehensive analysis of m^6^A methylation regulated by ALKBH5 (related to Figure 7).**
Total RNAs were extracted from PC9 cells transfected with siALKBH5#1, siALKBH5#3, or siNC (n = 3 for each group) and used for epitranscriptomic microarray analysis. (**a**) The m^6^A methylation level (%), which was calculated as immunoprecipitated transcript (IP)/all transcript (IP + Sup), of the nine samples were combined and plotted. (**b**) Correlations between the m^6^A methylation level and RNA length of each transcript were evaluated using Spearman’s correlation coefficient (r = 0.35). (**c**) Correlations between the m^6^A methylation level at baseline (cells transfected with siNC) and the difference of the m^6^A methylation level (%) following ALKBH5 knockdown [calculated as m^6^A in siALKBH5 − m^6^A in siNC (%)] were evaluated using Spearman’s correlation coefficient (r = −0.35). (**d** and **e**) Volcano plot showing differentially m^6^A-methylated RNAs in cells transfected with siALKBH5#1 (d) and siALKBH5#3 (e) compared with those using siNC. Dots indicate genes meeting both thresholds: *P* < 0.01 via the unpaired *t*-test and fold change of the m^6^A methylation level of >1.5 (log2 fold change = 0.5849, pink, hypermethylated) or of <0.67 (log2 fold change = −0.5849, light blue, hypomethylated). (**f**) Venn diagram indicating the number of hypermethylated genes in cells transfected with siALKBH5#1 or siALKBH5#3. (**g**) Gene set enrichment analysis with hallmark gene sets was performed using the methylation profiles of PC9 cells transfected with siALKBH5#1 (in the upper panel) and siALKBH5#3 (in the lower panel) and compared with those with siNC.

**Additional file 6: Figure S6. m^6^A methylation level quantified via MeRIP-qPCR using m^6^A antibody (related to Figure 7).**
(**a**) PolyA-enriched RNA extracted from PC9 cells was fragmented with RNA fragmentation buffer across various incubation times from 40 s to 70 s. The distribution of the size of the polyA-enriched RNA was evaluated using Agilent 4200 TapeStation. (**b**) PolyA-enriched RNA extracted from HEK293 cells was immunoprecipitated using anti-m^6^A antibody or normal IgG. The m^6^A level was calculated from transcript abundance in input or MeRIP fraction quantified via qPCR using primers for the positive control region [stop codon, (EEF1A1+)] or negative control region [exon 5, (EEF1A1−)] of human EEF1A1 (n = 3). (**c**) (Upper panel) Prediction scores of m^6^A modification in the MFAP5 gene were calculated using the SRAMP algorithm. The combined scores were distributed through the full-length mRNA as different levels of very high, high, moderate, and low confidence. Arrows show the location of qPCR primers. Adenosines in consensus sequences for m^6^A modification are presented in red. (Lower panel) PolyA-enriched RNA extracted from PC9 cells transfected with siALKBH5#1, siALKBH5#3, or siNC (n = 3) was immunoprecipitated using anti-m^6^A antibody or normal IgG. The m^6^A level was calculated from transcript abundance in input or MeRIP fraction quantified using qPCR. (**d**) MeRIP-qPCR for GADD45A and CASP14 was performed as described in (c). Results are presented as mean ± SD. **P* < 0.05, ***P* < 0.01, ****P* < 0.001, *****P* < 0.0001 indicates a significant difference between the indicated groups.

**Additional file 7: Figure S7. IGF2BPs knockdown efficacy and expression of CDKN1A and TIMP3 in the TCGA cohort (Related to Figure 8).**
(**a–d**) Relative mRNA expression levels of IGF2BPs (a–c) and ALKBH5 (d) were detected via qPCR in PC9 cells transfected with one of the siIGF2BPs and/or siALKBH5. Gene expression was normalized to the expression of GAPDH and is shown relative to those using siNC (n = 3). Results are presented as mean ± SD. (**e** and **f**) mRNA expression level of CDKN1A (e) and TIMP3 (f) in the paired non-cancerous and cancerous tissue of NSCLC in the TCGA database (n = 109 for each group). Bars indicate the median. *****P* < 0.0001 indicates a significant difference between the indicated groups.

**Additional file 8: Table S1. Primer list for qPCR**

**Additional file 9: Table S2. Clinical characteristics based on ALKBH5 and FTO expression in non-small cell lung cancer**

Differences between experimental groups were assessed using Mann–Whitney *U* test for continuous variables or Fisher’s exact test for categorical data. Data are presented as median (range) or number (n) (%).

**Additional file 10: Table S3. Multivariate Cox hazards models of survivals in all patients with non-small cell lung cancer**

Prediction of mortality of patients with non-small lung cancer. The univariate and multivariate Cox proportional hazards models were applied to generate the hazard ratios (HRs) of death. Multivariate analysis was adjusted by age, sex, smoking status, histology, stage, and ALKBH5.

**Additional file 11: Table S4. Differentially upregulated genes in expression microarray analysis**
The thresholds for differentially upregulated genes were set to fold change of >1.5 and *P* < 0.01 for Student’s *t*-test.

**Additional file 12: Table S5. Differentially downregulated genes in expression microarray analysis**
The thresholds for differentially downregulated genes were set to fold change of <0.67 and *P* < 0.01 for Student’s *t*-test.

**Additional file 13: Table S6. Expression of genes associated with m^6^A modification enzymes in expression microarray analysis**

**Additional file 14: Table S7. Hypermethylated genes in epitranscriptomic microarray analysis**
The thresholds for hypermethylated genes were set to fold change of >1.5 and *P* < 0.05 for unpaired *t*-test.
