## Supplementary Figures for "m^6^A demethylase ALKBH5 promotes tumor cell proliferation by destabilizing IGF2BPs target genes and worsens the prognosis of patients with non-small cell lung cancer"

Figure S1

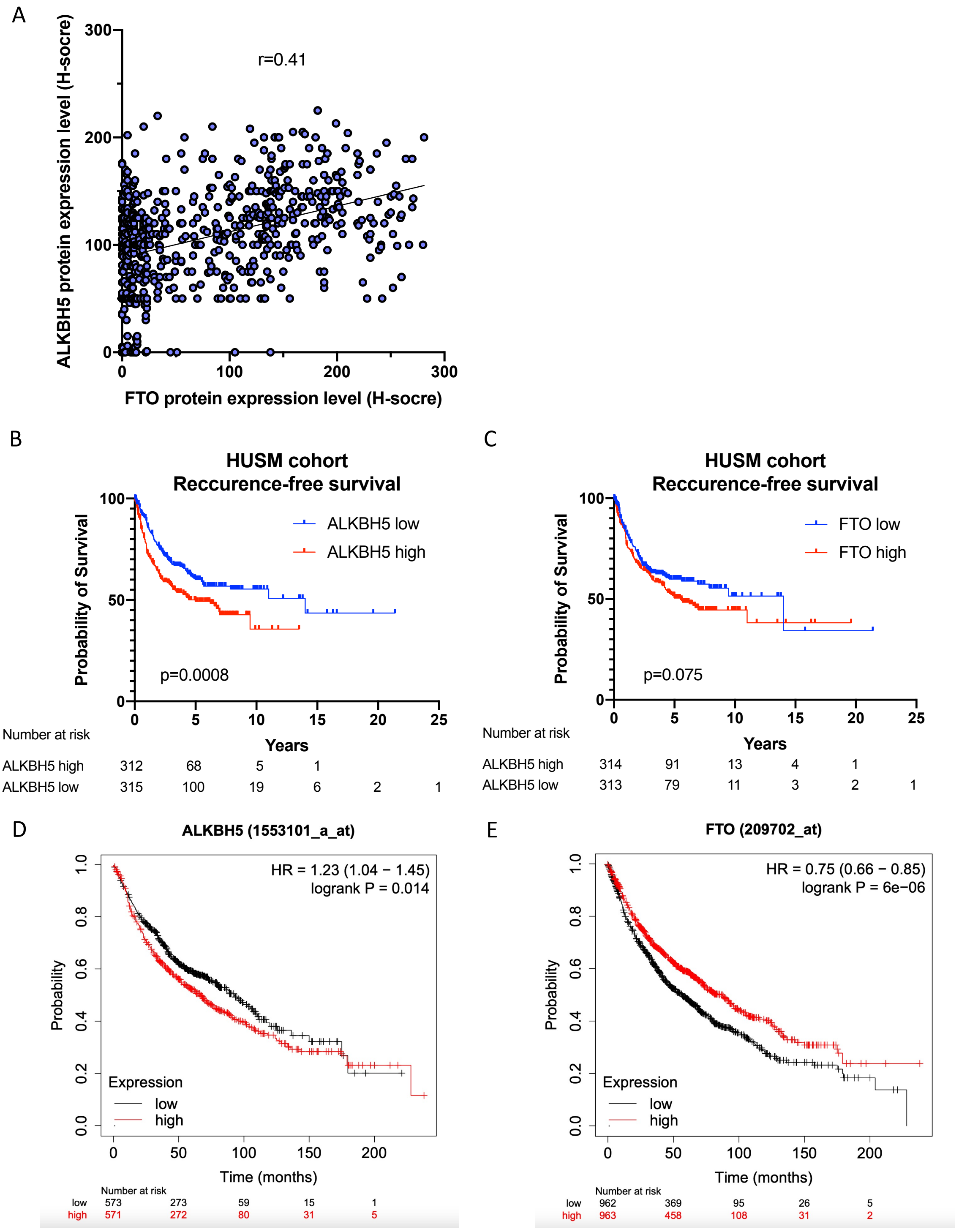
A
r=0.41
ALKBH5 protein expression level (H-socre)
FTO protein expression level (H-socre)
0
100
200
300
B
HUSM cohort
Reccurence-free survival
ALKBH5 low
ALKBH5 high
Probability of Survival
p=0.0008
Years
Number at risk
ALKBH5 high 312 68 5 1
ALKBH5 low 315 100 19 6 2 1
C
HUSM cohort
Reccurence-free survival
FTO low
FTO high
Probability of Survival
p=0.075
Years
Number at risk
ALKBH5 high 314 91 13 4 1
ALKBH5 low 313 79 11 3 2 1
D
ALKBH5 (1553101_a_at)
HR = 1.23 (1.04 − 1.45)
logrank P = 0.014
Probability
Expression
low
high
Time (months)
Number at risk
low 573 273 59 15 1
high 571 272 80 31 5
E
FTO (209702_at)
HR = 0.75 (0.66 − 0.85)
logrank P = 6e−06
Probability
Expression
low
high
Time (months)
Number at risk
low 962 369 95 26 5
high 963 458 108 31 2

Figure S2

A

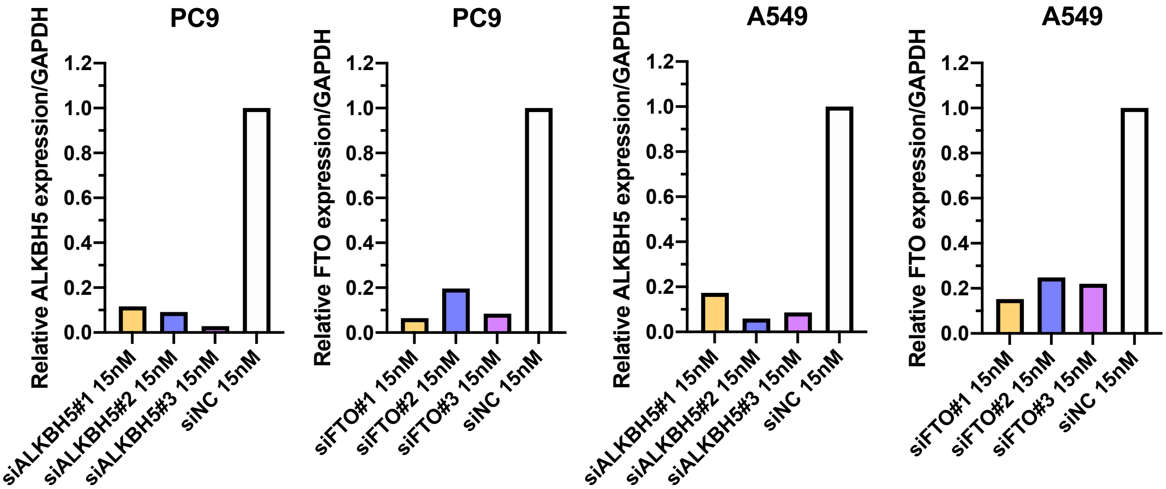

PC9
Relative ALKBH5 expression/GAPDH
1.2
1.0
0.8
0.6
0.4
0.2
0.0
siALKBH5#1 15nM
siALKBH5#2 15nM
siALKBH5#3 15nM
siNC 15nM
PC9
Relative FTO expression/GAPDH
siFTO#1 15nM
siFTO#2 15nM
siFTO#3 15nM
siNC 15nM
A549
Relative ALKBH5 expression/GAPDH
A549
Relative FTO expression/GAPDH

Figure S3

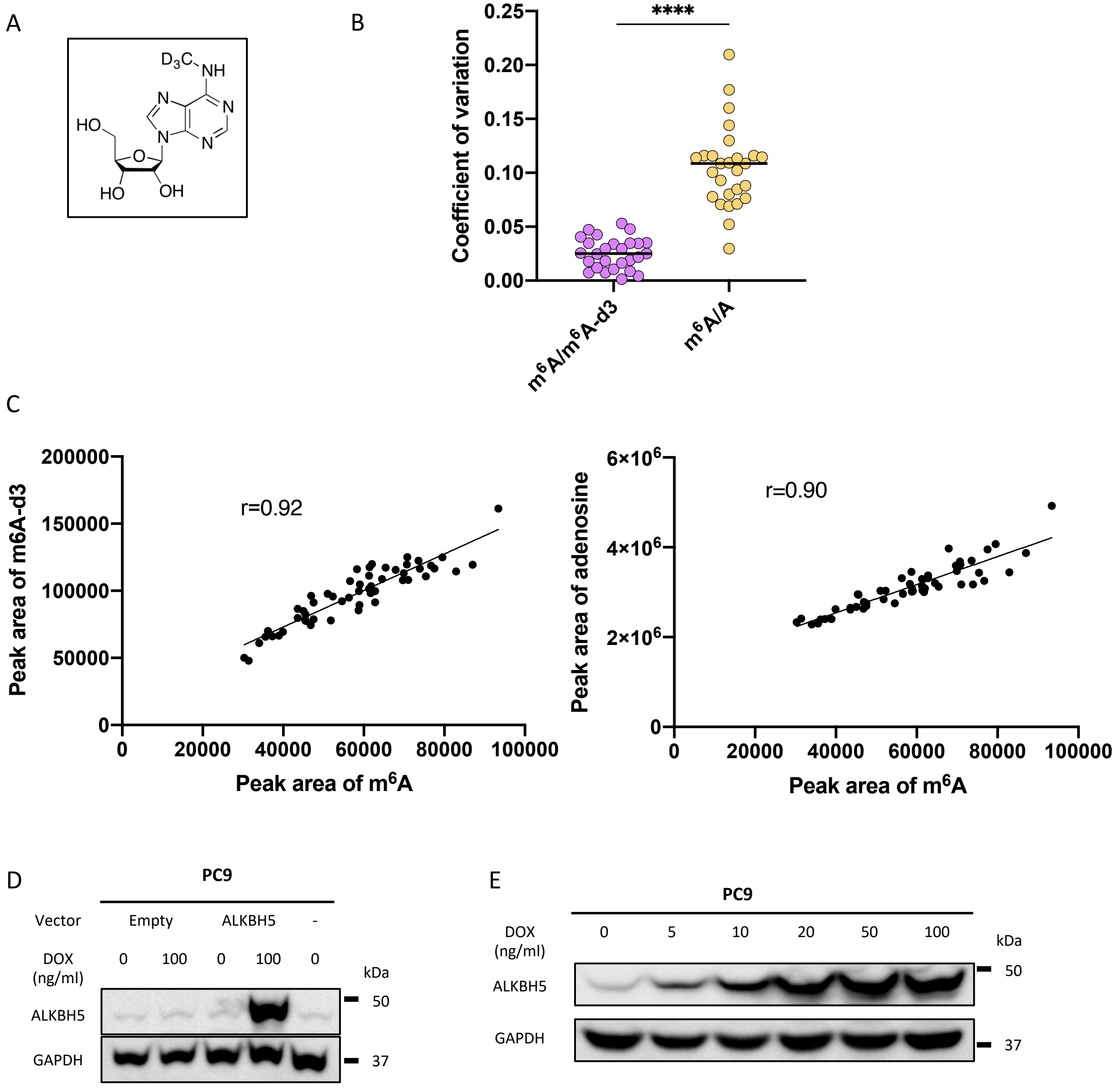
A
D3C
NH
N
N
N
N
HO
O
HO
OH
B
Coefficient of variation
0.25
0.20
0.15
0.10
0.05
0.00
****
m6A/m6A-d3
m6A/A
C
Peak area of m6A-d3
200000
150000
100000
50000
0
r=0.92
0
20000
40000
60000
80000
100000
Peak area of m6A
Peak area of adenosine
6×10^6
4×10^6
2×10^6
0
r=0.90
0
20000
40000
60000
80000
100000
Peak area of m6A
D
PC9
Vector
Empty
ALKBH5
-
DOX
(ng/ml)
0
100
0
100
0
kDa
ALKBH5
50
GAPDH
37
E
PC9
DOX
(ng/ml)
0
5
10
20
50
100
kDa
ALKBH5
50
GAPDH
37

### Figure S4

A

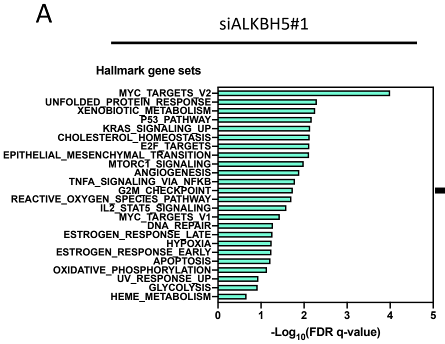


**Common enrichment gene sets between siALKBH5#1 and siALKBH5#3**

ANGIOGENESIS
APOPTOSIS
EPITHELIAL_MESENCHYMAL_TRANSITION
ESTROGEN_RESPONSE_EARLY
G2M_CHECKPOINT
HYPOXIA
KRAS_SIGNALING_UP
MYC_TARGETS_V2
P53_PATHWAY
TNFA_SIGNALING_VIA_NFKB
UNFOLDED_PROTEIN_RESPONSE
UV_RESPONSE_UP

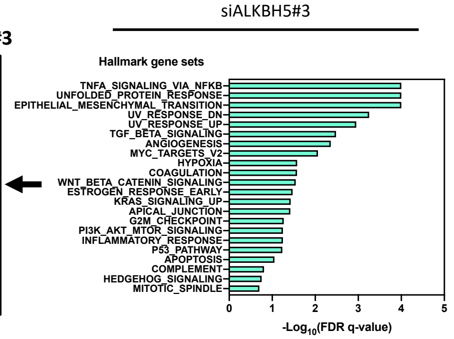


B

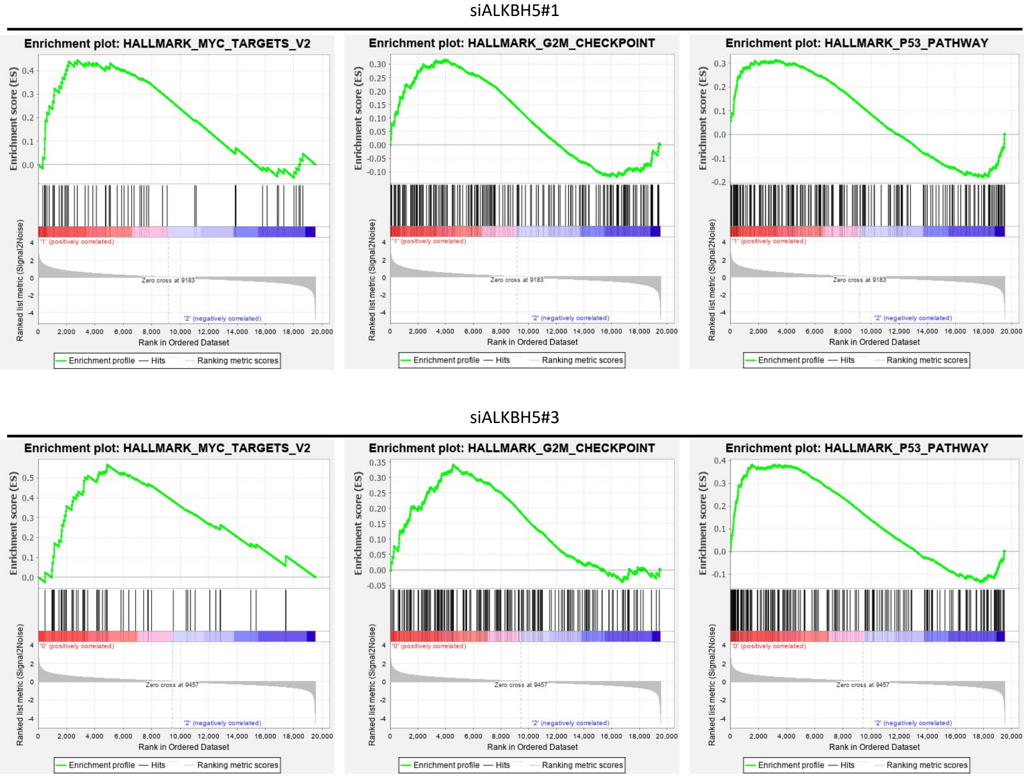

Figure S5

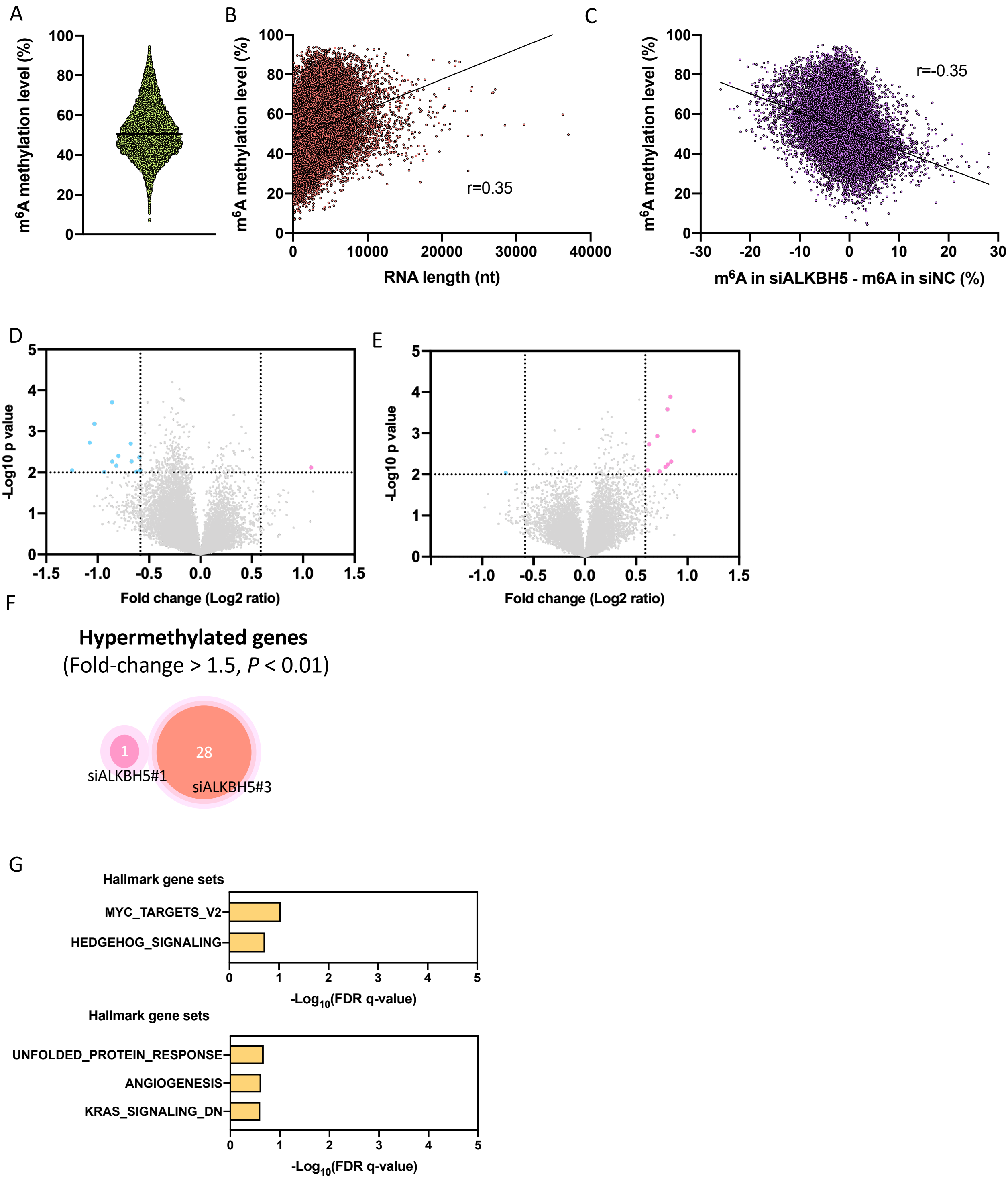
A
m6A methylation level (%)
100
80
60
40
20
0
B
m6A methylation level (%)
r=0.35
RNA length (nt)
0
10000
20000
30000
40000
C
m6A methylation level (%)
r=-0.35
m6A in siALKBH5 - m6A in siNC (%)
-30
-20
-10
0
10
20
30
D
-Log10 p value
Fold change (Log2 ratio)
-1.5
-1.0
-0.5
0.0
0.5
1.0
1.5
E
-Log10 p value
Fold change (Log2 ratio)
F
Hypermethylated genes
(Fold-change > 1.5, P < 0.01)
1
28
siALKBH5#1
siALKBH5#3
G
Hallmark gene sets
MYC_TARGETS_V2
HEDGEHOG_SIGNALING
-Log10(FDR q-value)
Hallmark gene sets
UNFOLDED_PROTEIN_RESPONSE
ANGIOGENESIS
KRAS_SIGNALING_DN
-Log10(FDR q-value)

Figure S6

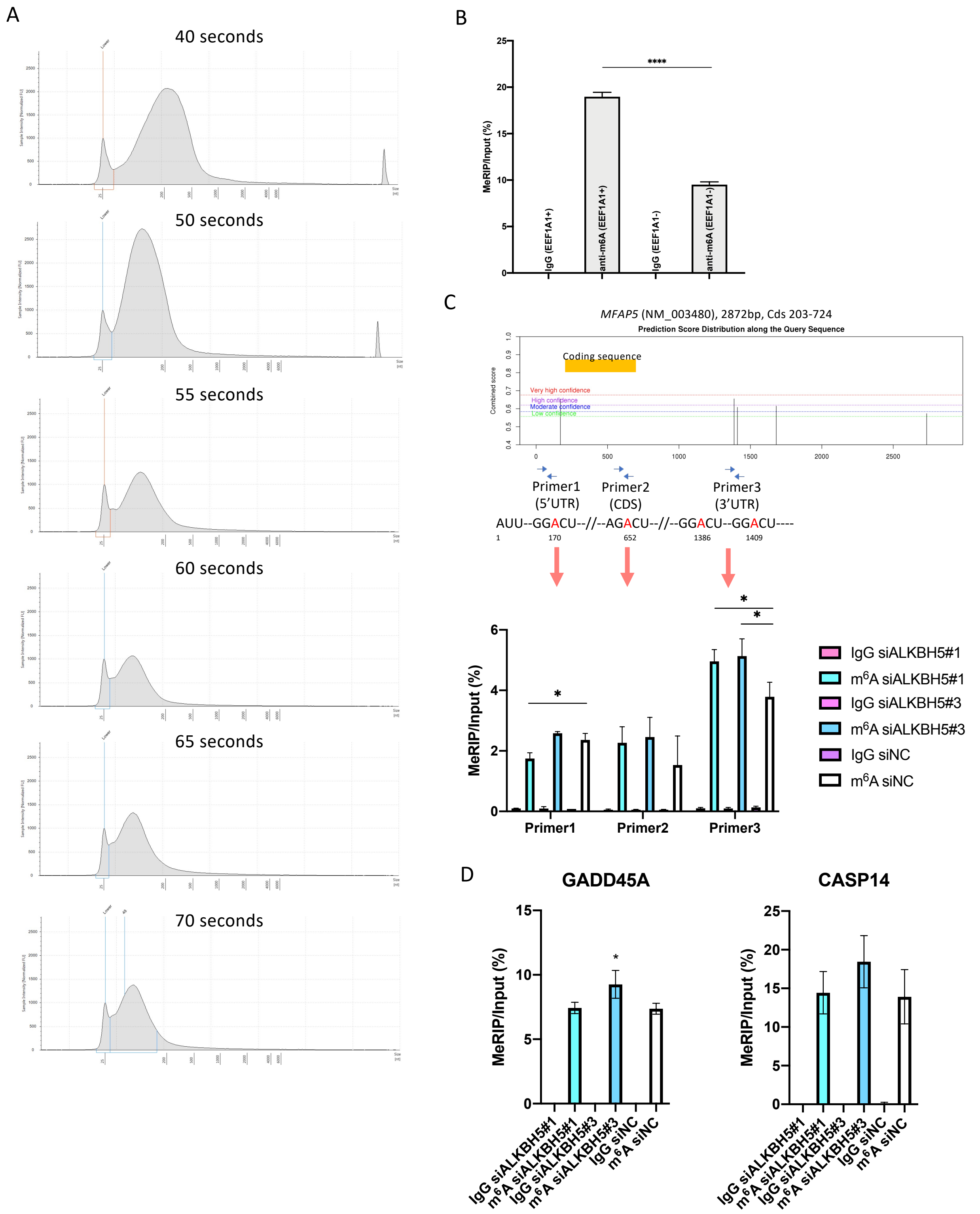
A
40 seconds
50 seconds
55 seconds
60 seconds
65 seconds
70 seconds
B
MeRIP/Input (%)
****
IgG (EEF1A1+)
anti-m6A (EEF1A1+)
IgG (EEF1A1-)
anti-m6A (EEF1A1-)
C
MFAP5 (NM_003480), 2872bp, Cds 203-724
Prediction Score Distribution along the Query Sequence
Coding sequence
Very high confidence
High confidence
Moderate confidence
Low confidence
Combined score
Primer1 (5'UTR)
Primer2 (CDS)
Primer3 (3'UTR)
AUU--GGACU--//--AGACU--//--GGACU--GGACU----
1
170
652
1386
1409
MeRIP/Input (%)
IgG siALKBH5#1
m6A siALKBH5#1
IgG siALKBH5#3
m6A siALKBH5#3
IgG siNC
m6A siNC
D
GADD45A
CASP14
MeRIP/Input (%)

Figure S7

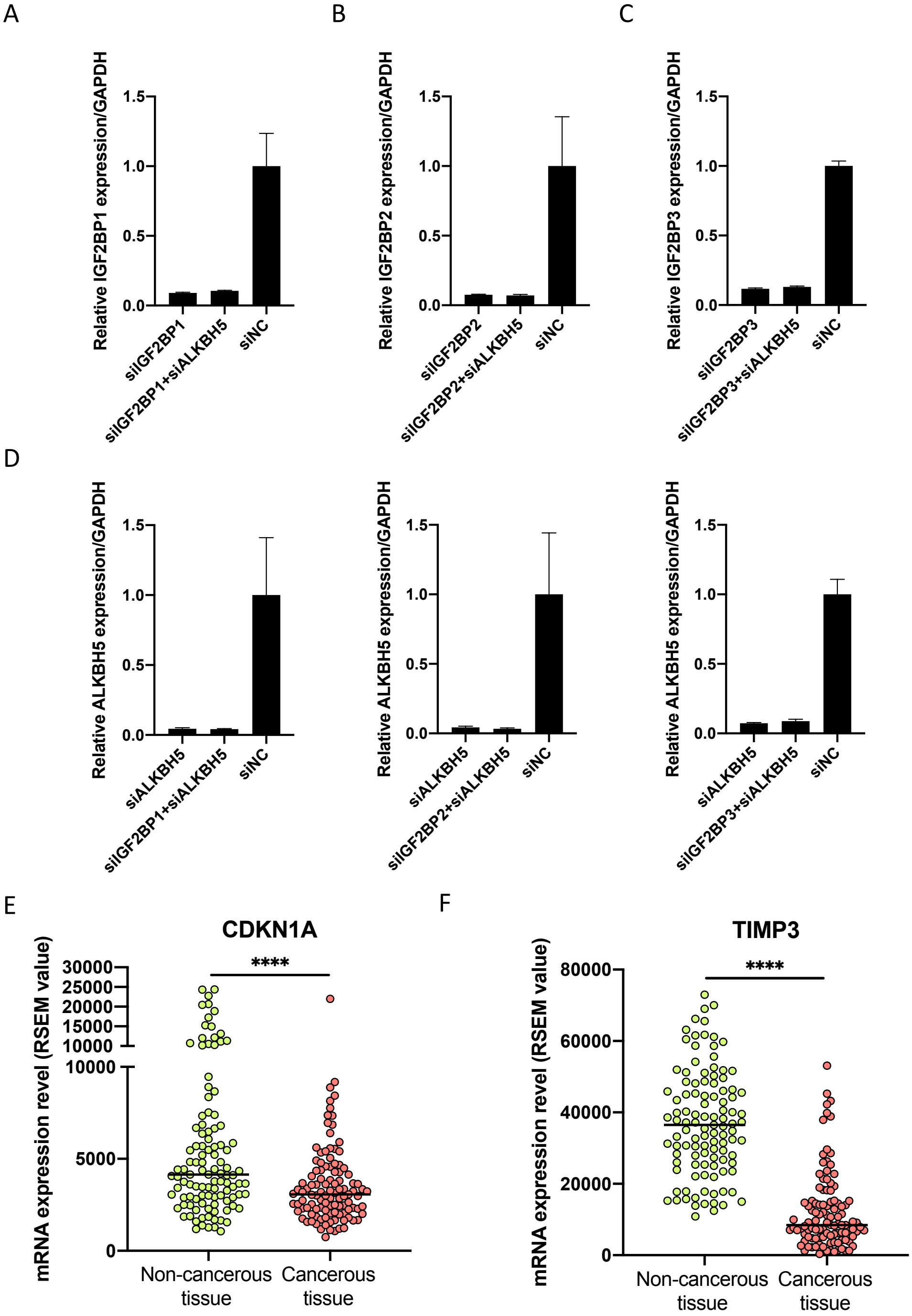

A
B
C
Relative IGF2BP1 expression/GAPDH
1.5
1.0
0.5
0.0
siIGF2BP1
siIGF2BP1+siALKBH5
siNC
Relative IGF2BP2 expression/GAPDH
siIGF2BP2
siIGF2BP2+siALKBH5
siNC
Relative IGF2BP3 expression/GAPDH
siIGF2BP3
siIGF2BP3+siALKBH5
siNC
D
Relative ALKBH5 expression/GAPDH
siALKBH5
siIGF2BP1+siALKBH5
siNC
siIGF2BP2+siALKBH5
siIGF2BP3+siALKBH5
E
CDKN1A
mRNA expression revel (RSEM value)
30000
25000
20000
15000
10000
10000
5000
0
****
Non-cancerous tissue
Cancerous tissue
F
TIMP3
80000
60000
40000
20000
0
